# Evolutionary dynamics of sex-biased genes expressed in cricket brains and gonads

**DOI:** 10.1101/2020.07.07.192039

**Authors:** Carrie A. Whittle, Arpita Kulkarni, Cassandra G. Extavour

**Affiliations:** Department of Organismic and Evolutionary Biology, Harvard University, 16 Divinity Avenue, Cambridge MA 02138, USA; Department of Molecular and Cellular Biology, Harvard University, 16 Divinity Avenue, Cambridge MA 02138, USA

**Keywords:** *Gryllus*, sex-biased expression, dN/dS, pleiotropy, gonad, brain, mating biology

## Abstract

Sex-biased gene expression, particularly sex-biased expression in the gonad, has been linked to rates of protein sequence evolution (nonsynonymous to synonymous substitutions, dN/dS) in animals. However, in insects, sex-biased expression studies remain centered on a few holometabolous species, and moreover, other major tissue types such as the brain remain underexplored. Here, we studied sex-biased gene expression and protein evolution in a hemimetabolous insect, the cricket *Gryllus bimaculatus*. We generated novel male and female RNA-seq data for two sexual tissue types, the gonad and somatic reproductive system, and for two core components of the nervous system, the brain and ventral nerve cord. From a genome-wide analysis, we report several core findings. Firstly, testis-biased genes had accelerated evolution, as compared to ovary-biased and unbiased genes, which was associated with positive selection events. Secondly, while sex-biased brain genes were much less common than for the gonad, they exhibited a striking tendency for rapid protein evolution, an effect that was stronger for the female than male brain. Further, some sex-biased brain genes were linked to sexual functions and mating behaviors, which we suggest may have accelerated their evolution via sexual selection. Thirdly, a tendency for narrow cross-tissue expression breadth, suggesting low pleiotropy, was observed for sex-biased brain genes, suggesting relaxed purifying selection, which we speculate may allow enhanced freedom to evolve adaptive protein functional changes. The findings of rapid evolution of testis-biased genes and male and female-biased brain genes are discussed with respect to pleiotropy, positive selection, and the mating biology of this cricket.

## 1 INTRODUCTION

Sexual dimorphism in animals is thought to be driven by differential gene expression, as most genes are common to both sexes (Ellegren & Parsch, 2007; Ingleby et al., 2014; Grath & Parsch, 2016). Sex-biased gene expression, and particularly male-biased gene expression, has been widely linked to rapid protein sequence evolution in studied animals (reviewed by (Ellegren & Parsch, 2007; Ingleby et al., 2014; Grath & Parsch, 2016)). In the insects, studies have largely focused on the holometabolous insect *Drosophila*, and have repeatedly shown the rapid evolution (high nonsynonymous to synonymous substitution rates, dN/dS) of male-biased genes, particularly those from the male sex cells or gonads, as compared to their female counterparts and/or to sexually unbiased genes (Jagadeeshan & Singh, 2005; Ellegren & Parsch, 2007; Haerty et al., 2007; Zhang et al., 2007; Jiang & Machado, 2009; Meisel, 2011; Grath & Parsch, 2012; Perry et al., 2015; Whittle & Extavour, 2019) (but see also (Dorus et al., 2006)). This pattern was also recently observed for the gonads of red flour beetles (*T. castaneum*) (Whittle et al., 2020). The rapid divergence of male-biased genes has been proposed to be due to adaptive changes in amino acids arising from sexual selection pressures including male-male and sperm competition (Swanson et al., 2001; Zhang et al., 2004; Proschel et al., 2006; Haerty et al., 2007), but could also reflect low pleiotropy that may relax purifying selection (Zhang et al., 2007; Mank & Ellegren, 2009; Assis et al., 2012; Dean & Mank, 2016; Whittle & Extavour, 2019). Nonetheless, the pattern of accelerated evolution of male-biased genes is not universal, as an opposite pattern of rapid evolution of female-biased, including ovary-biased, genes has been found in some holometabolous insects, namely mosquitoes (*Aedes*, *Anopheles*) (Papa et al., 2017; Whittle & Extavour, 2017). This difference from flies may reflect variation in their mating biology, whereby female-female competition for suitable males or male-mate choice may be more common in mosquitoes than in flies, and/or reflect variation in male- and female-related purifying selection among insects (Whittle & Extavour, 2017). At present however, given the narrow scope of insects studied to date, further investigation of sex-biased expression in the reproductive system and protein evolution is warranted, particularly in models outside the Holometabola.

While studies of sex-biased expression and its link to protein sequence evolution have largely focused on the reproductive system, a major, and markedly understudied structure, in terms of molecular evolution, is the brain. The brain is a major tissue type providing the neurological basis for the mating behaviors of courtship, intrasex competition, mate-choice, and post-mating male-female responses (Mank et al., 2007; Dalton et al., 2010; Naurin et al., 2011; Wright & Mank, 2013). Male and female differences in gene expression *per se* in the brain have been examined in some insects and vertebrates (Jagadeeshan & Singh, 2005; Mank et al., 2007; Santos et al., 2008; Small et al., 2009; Naurin et al., 2011; Catalan et al., 2012; Wright & Mank, 2013; Ingleby et al., 2014; Tomchaney et al., 2014; Huylmans & Parsch, 2015; Shi et al., 2016; Yang et al., 2016; Khodursky et al., 2020). Further, in *Drosophila*, analyses of a small number of neural genes showed a direct connection to mating functions and behaviors (Drapeau et al., 2003; Kadener et al., 2006; Dauwalder, 2008). However, there is a striking paucity of data on whether and how sex-biased expression in the brain is associated with protein sequence evolution (Mank et al., 2007; Wright & Mank, 2013). Moreover, the minimal research available from birds, humans and flies has suggested that male and female expression may have different effects on the rates of protein evolution, depending on the system (Mank et al., 2007; Shi et al., 2016; Khodursky et al., 2020) (see also some brain-related (Biswas et al., 2016) and composite-tissue analyses (Catalan et al., 2018; Congrains et al., 2018)), and the causes of those patterns remain poorly understood. It is therefore evident that additional study of sex-biased expression in the brain is needed, particularly with respect to its relationship to molecular evolution.

An insect model system that offers significant opportunities to address these problems is the cricket *Gryllus* (Order Orthoptera). *Gryllus* is a hemimetabolous insect, and thus in an outgroup order to the Holometabola (Misof et al., 2014). The two-spotted cricket *G. bimaculatus* in particular has emerged as a significant insect model in biology, including for genetics, neuroscience and germ line establishment and development (Kulkarni & Extavour, 2019). In fact, many of the developmental mechanisms of *G. bimaculatus* appear more typical of arthropods than the widely studied, and relatively derived, model *Drosophila melanogaster* (Mito & Noji, 2008; Donoughe & Extavour, 2016). Moreover, many aspects of its mating biology are currently well understood. *G. bimaculatus* exhibits intense male-male and sperm competition, including aggressive male-male fighting and mate guarding (Vedenina & Shestakov, 2018; Gee, 2019), increased rates of male transfer of spermatophores to females in the presence of other males (Lyons & Barnard, 2006), and the complete mixing of sperm from multiple males in the storage organ of the female reproductive tract, the spermatheca (Simmons, 1986; Morrow & Gage, 2001). In addition, females have shown preferences for novel and young mating partners (Zhemchuzhnikov et al., 2017), and for males with larger body size and higher quality auditory signals (Bateman et al., 2001; Zhemchuzhnikov et al., 2017). Females also exhibit a post-mating behaviour of removing spermatophores of non-favored males from their reproductive tract (Simmons, 1986), suggesting a propensity for female mate choice in this organism. Moreover, in terms of the brain, experiments in *G. bimaculatus* have shown that the brain is directly involved in male mating behaviors such as courtship, copulation, spermatophore protrusion, mating intervals and male-female auditory mating signalling (Matsumoto & Sakai, 2000; Haberkern & Hedwig, 2016; Sakai et al., 2017). The study of *Gryllus* therefore provides a valuable avenue to advance our knowledge of sex-biased expression in reproductive and brain tissues, including relationships to dN/dS and pleiotropy, in a taxon having well-studied mating biology.

Here, we assess sex-biased gene expression for two tissue types from the reproductive system (gonad and somatic reproductive system) and from the nervous system (brain and ventral nerve cord) in *G. bimaculatus,* and evaluate their relationships to protein sequence evolution. We report that male-biased gene expression in the gonad is linked to rapid protein sequence evolution (dN/dS), as compared to unbiased and female-biased genes. However, we observed no consistent effect of sex-biased expression in the somatic reproductive system (non-germ line tissues) on dN/dS, despite the roles of these sexual tissues in male-female interaction, mating and fertilization, and their potential exposure to sexual selection pressures (Swanson & Vacquier, 2002; Swanson et al., 2004; Clark & Swanson, 2005; Panhuis & Swanson, 2006; Haerty et al., 2007). With respect to the brain, we demonstrate that sex-biased genes are uncommon as compared to the gonad, and that these genes typically evolve very rapidly, especially the female-biased brain genes. Further, sex-biased brain genes are conspicuously linked to predicted sex-related functions. The sex-biased brain genes exhibit especially low cross-tissue expression, a proxy for pleiotropy (Mank & Ellegren, 2009), which may in itself accelerate protein sequence evolution due to relaxed purifying constraint. We propose that this low pleiotropy may also comprise a mechanism potentially allowing greater freedom for these brain-expressed proteins to evolve adaptive functional changes, an evolutionary dynamic that has been suggested in some studies (Otto, 2004; Larracuente et al., 2008; Mank et al., 2008; Mank & Ellegren, 2009; Meisel, 2011). We consider the putative roles of the male and female mating biology of *G. bimaculatus* in shaping the present findings.

## 2 MATERIALS AND METHODS

### 2.1 Biological samples and RNA-seq

For our RNA-seq assessment of *G. bimaculatus* we isolated the male and female gonad, somatic reproductive system, brain and ventral nerve cord (shown in Fig. 1, Table S1; schematic is based on (Kumashiro & Sakai, 2001) and simplified from Fox 2001; http://lanwebs.lander.edu/faculty/rsfox/invertebrates/acheta.html)). RNA-seq data were obtained for four paired male and female tissue types from adult virgins (biological replicates (Congrains et al., 2018) and read counts in Table S1). The somatic (non-germ line related) reproductive system herein for males included the pooled vasa deferentia, seminal vesicles and ejaculatory duct, and for females included the spermatheca, common and lateral oviducts, and bursa (Fig. 1A,B). A ninth, unpaired reproductive tissue type, the male accessory gland (Fig. 1G), was also extracted, as its gene expression has been linked to protein sequence changes (Swanson et al., 2001; Clark & Swanson, 2005; Haerty et al., 2007), and it provides an additional sexual tissue type for the analysis of cross-tissue expression breadth (see below in Methods). Further, we considered that its inclusion in the male somatic reproductive system sample might overwhelm the transcript population of that tissue type upon RNA-seq, and make it incomparable to the female reproductive system.

**Fig. 1.**
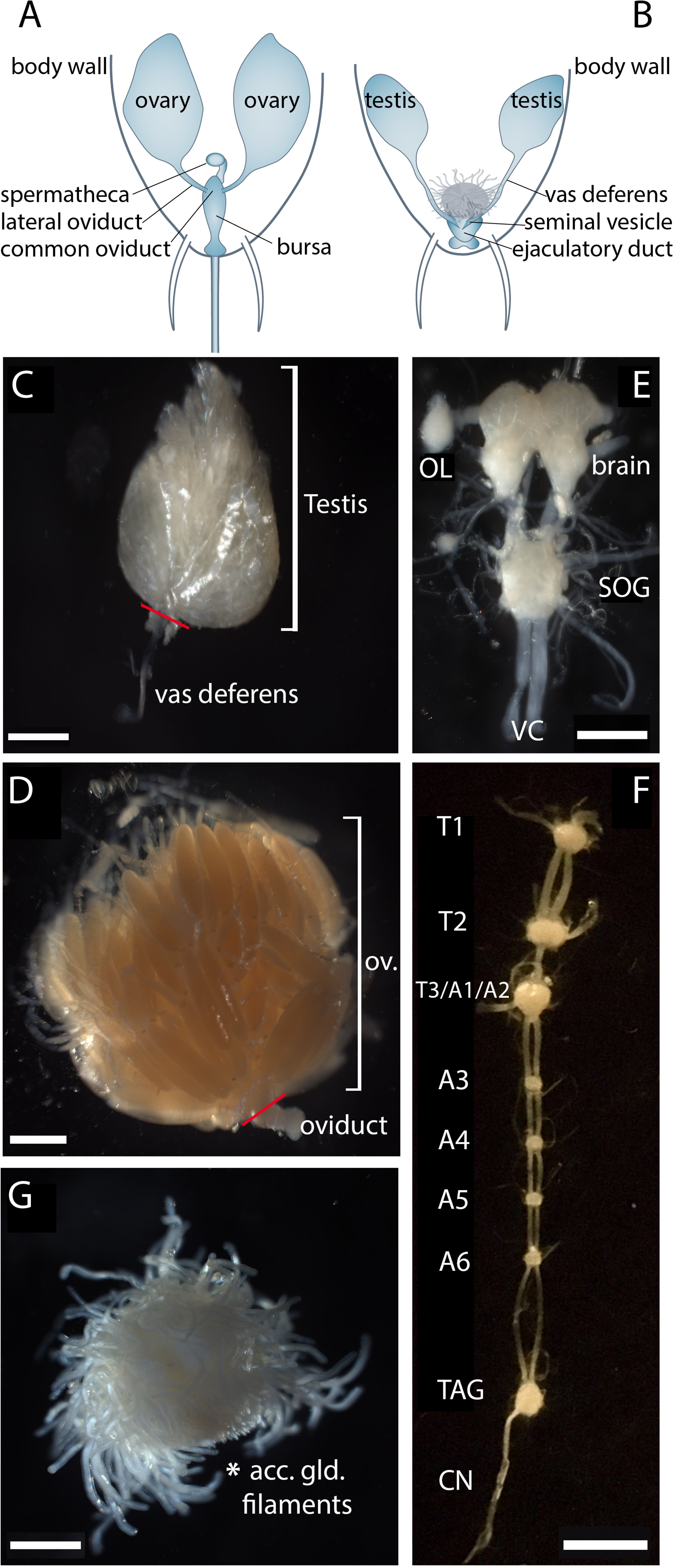
*Gryllus bimaculatus* reproductive and nervous system tissues studied herein. A) Schematic diagram of the female reproductive system showing the gonads and the somatic tissues included in the somatic reproductive system under study. B) Schematic diagram for the male gonads and male somatic reproductive system. C-G provide micrographs of various tissue types studied herein. C) the testis (one testis shown here; both testes from a given male were used for sampling), including a part of its attached vas deferens (boundary indicated by red line; the vasa deferentia were included in the male somatic reproductive system libraries, and not in testis libraries). D) the ovary (ov; one ovary shown here; both ovaries from a given female were used for sampling) and an immediately attached segment of oviduct (boundary indicated by red line; the oviducts were included in the female somatic reproductive system libraries, and not in ovary libraries). E) the brain, including an optic lobe (OL) (one OL shown here; both OLs from a given individual were included in brain samples). For context, the attached suboesophageal ganglion (SOG) and upper portion of the ventral nerve cord (VC) are also shown; these structures were included in the ventral nerve cord libraries and not in brain libraries. F) the ventral nerve cord including the three thoracic ganglia (T1: prothoracic, T2: mesothoracic, T3/A1/A2: metathoracic ganglion complex), and five abdominal ganglia (A3-A6 and the terminal abdominal ganglion TAG) (Huber, 1963; Jacob & Hedwig, 2016). The cercal nerve (CN) of one side is also shown. For the ventral nerve cord samples, all tissues in F and the SOG were pooled. G) The male accessory gland consisting of numerous accessory gland filaments (asterisk; also shown schematically as filamentous structure in B). Scale bars: 500μm in C and E, 1000μm in D and G, 2500μm in F.

The rearing of specimens for tissue sampling was as follows: post hatching, wild type *G. bimaculatus* nymphs from an existing laboratory colony inbred for at least 14 years were grown at 29°C until adulthood in well-ventilated plastic cages on a 12 hour day/ 12 hour night cycle (Kainz et al., 2011). Plastic cages were provided with egg cartons for shelter, and the animals were fed with ground cat food (Purina item model number 178046) and water. Prior to the final nymphal molt, animals were sexed based on the presence (female) or absence (male) of an ovipositor and separated into male and female cages to avoid any mating and thus obtain virgin samples. Dissections were then performed on the unmated adults within a week after their final molt, by briefly anesthetizing the animals on ice for 5-10 minutes prior to dissection. Different tissue types (gonad, somatic reproductive system, brain, ventral nerve cord, male accessory reproductive glands) were dissected per animal using sterile equipment wiped with ethanol and RNaseZap (Ambion, catalog number AM9780), in ice-cold 1x Phosphate Buffer Saline (PBS), and the tissue cleaned of any unwanted contaminating material. Each tissue was then transferred immediately into individual 1.5ml Eppendorf tubes containing 500μl of pre-frozen Trizol (Thermo Fisher, catalog number 15596018) on dry ice, and stored at −80°C until further use. RNA extractions, library processing and RNA-seq were then performed as described previously (Whittle et al., 2020). The same procedure was conducted for specimens of *G. assimilis*, which was used to obtain RNA-seq data for an assembly to be used for dN/dS analysis (Table S2; which also included a carcass tissue type). The *G. assimilis* eggs were obtained from the Hedwig lab (University of Cambridge, UK) and reared to adulthood, using the same animal husbandry protocols as published previously for *G. bimaculatus* (Mito & Noji, 2008; Kainz et al., 2011; Kochi et al., 2016).

The RNA-seq reads (76bp in length) for each sample were trimmed of adapters and poor quality bases using the program BBduk available from the Joint Genome Institute (https://jgi.doe.gov/data-and-tools/bbtools/) using default parameters.

### 2.2 CDS of *G. bimaculatus* and sex-biased gene expression

The CDS of our main target species *G. bimaculatus* were obtained from its recently available genome (Ylla et al., 2021). The annotated genome had 17,714 predicted transcripts (after selecting the longest CDS per gene; (Ylla et al., 2021)). For this gene set, we extracted the CDS with a start codon, no ambiguous nucleotides, and at least 150bp in length, yielding 15,539 CDS for study (mean length=417.0 codons/CDS ±3.5 (standard error (SE))) for *G. bimaculatus*. For analysis of sex-biased gene expression in *G. bimaculatus*, the expression level for each of 15,539 *G. bimaculatus* genes was determined by mapping reads from each RNA-seq dataset per tissue to the full CDS list using Geneious Read Mapper (Kearse et al., 2012), a program previously found to be as effective as other common read mappers (cf. Whittle et al., 2020). We compared expression between males and females for the gonad, somatic reproductive system, brain, and ventral nerve cord using the program DESeq2, which uses the mapped reads across biological replicates and the negative binomial distribution to quantify the P-values of expression differences (Love et al., 2014). In addition, the degree of sex-biased expression per gene was determined using the ratio of average in FPKM across the replicates for female and male tissues. Any gene that had a two-fold or greater ratio in average expression in one sex (as compared to the other) and a statistically significant P-value from DESeq (P<0.05) as well as a FPKM of at least 1 in one tissue type was defined as sex-biased (*cf.* on a two-fold cutoff (Proschel et al., 2006; Assis et al., 2012; Whittle & Extavour, 2017; Parker et al., 2019; Whittle & Extavour, 2019; Whittle et al., 2020). Given the use of two biological replicates for the large-scale RNA-seq (Table S1) and our high threshold cutoff (two-fold), the identification of sex-biased genes herein is conservative. All genes not defined as sex-biased per tissue type were defined as unbiased (Zhang et al., 2010; Whittle & Extavour, 2017; Darolti et al., 2018; Parker et al., 2019; Whittle & Extavour, 2019; Whittle et al., 2020), which herein includes all genes with less than two-fold sex-biased expression or with <1 FPKM (including undetectable expression, and apt not to play sex-related roles) in both females and males (Whittle & Extavour, 2017; Whittle & Extavour, 2019; Whittle et al., 2020). Thus, all 15,539 genes belonged to one of these three categories per tissue (Fig. 2). We note that 95.4% of the 15,539 *G. bimaculatus* genes were expressed in at least one of the nine tissues (Table S1), suggesting the vast majority of genes have putative roles in some or all of these studied tissues.

**Fig. 2.**
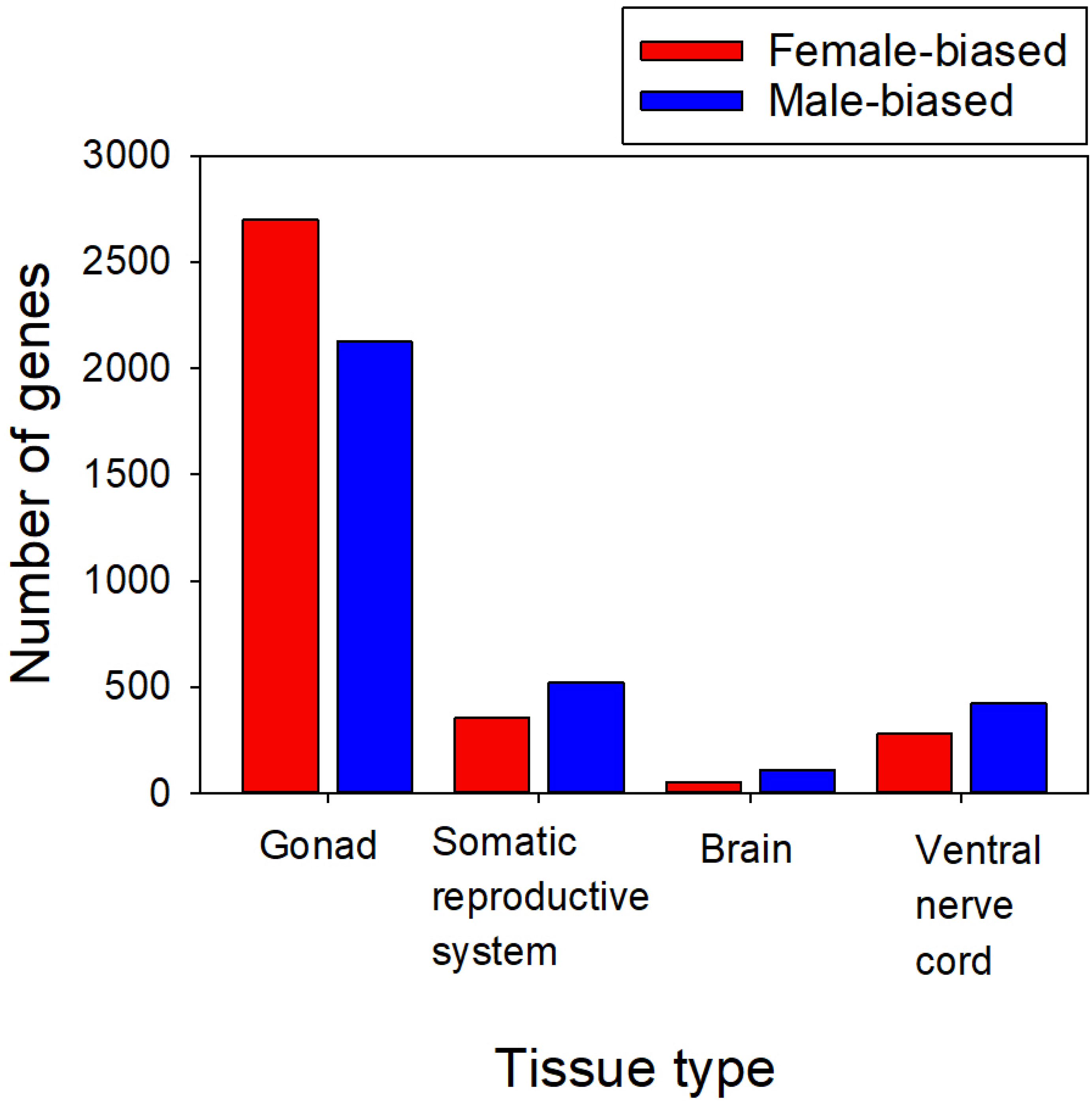
The number of male-biased and female-biased genes identified in the gonad, somatic reproductive system, brain, and ventral nerve cord across all 15,539 *G. bimaculatus* genes under study (sex-biased indicates a two-fold difference in expression and P<0.05). All remaining genes not shown per tissue type had unbiased status as follows: gonad (N=10,717), somatic reproductive system (N=14,666), brain (N=15,382) and ventral nerve cord (N=14,835).

### 2.3 Assembly of *G. assimilis* RNA-seq data and protein sequence divergence analysis

#### 2.3.1 Assembly of reads

We aimed to assess whether and how evolutionary pressures on protein sequence divergence, measured as dN/dS, varied with sex-biased gene expression. Unlike *Drosophila, Gryllus* is currently an emerging model genus with limited genomic resources outside the recent *G. bimaculatus* genome (Ylla et al., 2021). Thus, to measure dN/dS, we generated and assembled novel RNA-seq data for its sister species *G. assimilis* to obtain a CDS list for that organism (Table S2). Two-species assessments of dN/dS have been repeatedly shown to be an effective means to study divergence of sex-biased genes (*cf.* (Mank et al., 2007; Baines et al., 2008; Meisel, 2011; Assis et al., 2012; Whittle & Extavour, 2017; Jaquiery et al., 2018) including for organisms with few available genomes, as is the case with *Gryllu*s.

For *G. assimilis* we assembled all the trimmed reads for the RNA-seq datasets of *G. assimilis* shown in Table S2. For this, the *G. assimilis* reads were *de novo* assembled into contigs using Trinity (Grabherr et al., 2011) set to default parameters using Galaxy (https://usegalaxy.org/). We then identified CDS using the PlantTribes pipeline tools (Wall et al., 2008). To assess the completeness of the assembled transcriptome, we used BUSCO 3.0.1 (Seppey et al., 2019) to reveal the percentage of the single-copy CDS that was observed in the standardized Arthropod conserved gene set, and as employed in gVolante ((Nishimura et al., 2017) https://gvolante.riken.jp/analysis.html). To refine the CDS for *G. assimilis* we then assessed each CDS in ORF predictor, using its downloadable Perl script (Min et al., 2005), to identify the highest quality reading frame per sequence. In ORF predictor, we used the option to include the best-hit (lowest e-value) BLASTX alignment (conducted in BLAST+ v2.7.1, https://blast.ncbi.nlm.nih.gov) (Altschul et al., 1997) of *G. assimilis* versus the reference *G. bimaculatus* protein database (i.e., its translated 15,539 CDS) to define reading frames, and retained all *G. assimilis* CDS that were at least 150bp long and had a start codon. Details of the *G. assimilis* assembly, including BUSCO scores (Seppey, Manni et al. 2019), and ORF predictions (Min et al., 2005) are provided in Text File S1.

It is worth noting that while paired-end reads have often been used for RNA-seq assembly, transcriptome assemblies from single-end reads have been successfully employed to obtain CDS (not requiring isoforms) as studied herein (Gongora-Castillo & Buell, 2013; Hibsh et al., 2015). Further to this point, single-end reads have even been applied for *de novo* assemblies in non-traditional model systems (Gongora-Castillo & Buell, 2013; Hibsh et al., 2015). Here, we have the additional advantage of a closely related reference genome to *G. assimilis*, namely *G. bimaculatus* (Ylla et al., 2021), to identify and confirm orthologs.

#### 2.3.2 Ortholog identification and dN/dS

Gene ortholog matches between *G. bimaculatus* and *G. assimilis* were identified using reciprocal BLASTX of the full CDS list between the two species in the program BLAST+ v2.7.1 (https://blast.ncbi.nlm.nih.gov) (Altschul et al., 1997). Genes having an identical best match sequence (lowest e-value) in both forward and reverse contrasts and e<10^-6^ were defined as putative orthologs. The identified orthologous gene sequences in *G. bimaculatus* and *G. assimilis* were aligned by codons using MUSCLE (Edgar, 2004) set to default parameters in the program Mega-CC v7 (Kumar et al., 2012) and gaps removed. Removal of divergent regions from alignments, despite some loss of sequence regions, improves quantification of protein divergence; thus, highly divergent segments were excluded using the program GBlocks v. 0.91b set at default parameters (Castresana, 2000; Talavera & Castresana, 2007).

Using the aligned *G. bimaculatus* and *G. assimilis* CDS, we employed yn00 of PAML using the Yang and Nielson 2000 substitution model, which comprises a maximum likelihood method that accounts for codon usage biases (Yang & Nielsen, 2000; Yang, 2007), to measure dN, dS, and dN/dS (Yang, 2007) (note that dN/dS measures using Yang and Neilson 2000 (Yang & Nielsen, 2000) were strongly correlated to those using other models; e.g., values from the Pamilo and Bianchi 1993 method (Pamilo & Bianchi, 1993) had Spearman’s R=0.95 P<2X10^-7^). Values of dN/dS reflect the standardized rate of protein sequence evolution (dN to dS), whereby values >1, =1, and <1 suggest a prevalent history of positive selection, neutral evolution and purifying selection respectively (Yang, 2007). However, even when <1 for gene-wide measures of dN/dS, elevated values suggest greater roles of positive selection and/or relaxed purifying selection (Swanson & Vacquier, 2002; Buschiazzo et al., 2012). Genes that were best matches by reciprocal BLASTX, and for which both values of dN and dS values were <1.5 (and thus were unsaturated (Castillo-Davis et al., 2004; Treangen & Rocha, 2011)), were defined as high confidence orthologs (N=7,220) between *G. bimaculatus* and *G. assimilis* for dN/dS analysis. The low dN values between these two cricket species, which were typically well below 1 as described in the Results and Discussion, should allow precise detection of orthologs, not only for single copy genes but also for duplicated genes, which can evolve rapidly (Demuth et al., 2006; Hahn et al., 2007). Overall, given the strict criteria we used for identification of high confidence orthologs, the paired alignments and dN, dS, and dN/dS measures herein are conservative.

### 2.4 Pleiotropy

We assessed the expression breadth across tissues for *G. bimaculatus* using nine tissues, the four paired female and male tissues and the male accessory glands (Table S1), as a proxy for pleiotropy, or multifunctionality of a gene (Otto, 2004; Larracuente et al., 2008; Mank et al., 2008; Mank & Ellegren, 2009; Meisel, 2011; Assis et al., 2012; Dean & Mank, 2016; Whittle & Extavour, 2017). Note that we choose a direct determination of expression breadth, rather than an index ((Duret & Mouchiroud, 2000; Haerty et al., 2007; Meisel, 2011), see also (Yanai et al., 2005).

### 2.5 Positive selection tests

In our core assessments of gene-wide dN/dS using paired contrasts of *G. bimaculatus* and *G. assimilis* from the same genus, any values >1 were interpreted as an indicator of a potential history of positive selection (Swanson et al., 2001; Torgerson et al., 2002; Nielsen et al., 2005; Clark et al., 2006; Yang, 2007; Hunt et al., 2011; Buschiazzo et al., 2012; Ghiselli et al., 2018). For analysis of genes with dN/dS >1, we included only those genes with both dN and dS>0.

In addition to this assessment, we examined positive selection at specific codon sites for the *G. bimaculatus* branch using branch-site analysis in codeml of PAML (Yang, 2007). As an outgroup species was required for this assessment, we used the recently available assembled and annotated *Laupala kohalensis* genome (Blankers et al., 2018; Ylla et al., 2021). Three-way orthologs between *G. bimaculatus*, *G. assimilis*, and *L. kohalensis* were identified using reciprocal BLASTX (e<10^-6^) among each of the three paired species contrasts (our criterion was that for each *G. bimaculatus*-*G. assimilis* paired ortholog, the same matching *L. kohalensis* CDS must be found using reciprocal BLASTX to *G. bimaculatus* CDS and to *G. assimilis* CDS). Genes were aligned by codons using all three-species CDS and filtered using GBlocks (Castresana, 2000; Talavera & Castresana, 2007) and gaps removed as described in “*2.3.2 Ortholog identification and dN/dS*” (note: alignments using this relatively distant outgroup were conducted independently of the paired *Gryllus* alignments). The phylogeny was ((*G. bimaculatus*, *G. assimilis*), *L. kohalensis*) and was unrooted for the PAML free-ratio analysis (Model=1, NSsites=0 in codeml) that was used to determine dN and dS per branch. Only those genes with dN<3, and with dS<3 (Mank et al., 2007) in the *L. kohalensis* branch were defined as high confidence orthologs and used for branch-site analysis (unlike the two-species contrasts within *Gryllus*, which were more closely related and had a cut-off of 1.5). For genes meeting these criteria, branch-site positive selection was assessed on the *G. bimaculatus* branch using Chi-square values for 2XΔln Likelihood (P<0.05) between models without (null hypothesis) and with (alternate hypothesis) positive selection (Model=2, NSsites=2, omega fixed versus estimated) as described in the PAML manual (Yang, 2007). We note that our stringent approach to defining three-way orthologs, and the distance of the outgroup, favors study of the more conservative portion of the genome for branch-site analysis. Further, some studies have suggested that branch-site analysis can lack sensitivity to detect functional changes (Nozawa et al., 2009; Toll-Riera et al., 2011), and/or may generate false positives (Nozawa et al., 2009; Wisotsky et al., 2020), the latter likely being sensitive to the stringency of alignment. We thus aimed to control this factor by our strict approach to this assessment (excluding genes with any signs of dN or dS saturation).

### 2.6 Sex-biased expression between *G. bimaculatus* and *G. assimilis*

As a supplementary analysis to our core assessment of sex-biased expression in our main target taxon *G. bimaculatus*, we also examined sex-biased transcription of genes in *G. assimilis*. For this, expression was determined using its assembled CDS list (described in Text file S1 and in the Results and Discussion) and the RNA-seq data (Table S2), as was described for *G. bimaculatus*. We focused on the gonads, which had the highest number of sex-biased genes among tissues in *G. bimaculatus* (see Results and Discussion). We assessed the correlation in expression for orthologs between the two species using Spearman’s ranked correlations. In turn, we determined those genes with conserved and variable sex-biased expression status in the gonads between species, and their relationships to dN/dS.

### 2.7 Gene ontology

Gene ontology (GO) was characterized using the tool DAVID (Huang da et al., 2009). For this, we identified orthologs to *G. bimaculatus* in the insect model *D. melanogaster*, which has the most well-studied insect genome to date (CDS v6.24 available from www.flybase.org (Gramates et al., 2017)), using BLASTX (https://blast.ncbi.nlm.nih.gov) (Altschul et al., 1997) and the best match (lowest e-value with cut off of e<10^-3^ of *D. melanogaster*). Single direction BLASTX (Altschul et al., 1997) with *G. bimaculatus* CDS as the query to the *D. melanogaster* protein database was used for these assessments (unlike for the more rigorous reciprocal BLASTX analysis used to identify orthologs between the two *Gryllus* species for dN/dS analysis), as we considered that the latter would be overly conservative between these insects from different orders for the purpose of functional characterization and analysis, and might prevent detection of putative paralogs in the crickets. *D. melanogaster* gene identifiers were input into DAVID (Huang da et al., 2009) to obtain gene putative GO functions and/or classifications. The *D. melanogaster* BLASTX searches were used solely for identification of putative orthologs to ascertain putative GO functions for our sex-biased and unbiased genes (and for putative SFP identification), and were not used for any dN/dS analysis, which was restricted to genes aligned within the crickets.

### 2.8 Seminal fluid proteins

As a complementary reproductive assessment in *G. bimaculatus*, we examined seminal fluid proteins (SFPs). We used *D. melanogaster* as our reference for SFP identification given this species has the most well-studied insect genome, transcriptome, and proteome to date, thus providing a more complete profile than the currently available smaller and likely partial SFP lists for crickets, which were obtained using transcriptomics and/or proteomics from reproductive tissues (Andres et al., 2013; Simmons et al., 2013). A recent proteome analysis of sexual structures in *D. melanogaster* confirmed functions for 134 SFPs (Sepil et al., 2019). Thus, we identified potential orthologs in *G. bimaculatus* to these SFPs in *D. melanogaster* (using single direction BLASTX as conducted for GO analysis) and assessed whether those genes had high confidence orthologs (and their dN/dS values) between *G. bimaculatus* and *G. assimilis*.

### 2.9 Data Availability

All RNA-seq data for *G. bimaculatus* and *G. assimilis* for this study described in Tables S1 and S2 are available at the Short Read Archive (SRA; https://www.ncbi.nlm.nih.gov/sra) under the project identifier PRJNA56413 (under species name and Study ID SRP220521). The studied genome data are publicly available as previously described for *G. bimaculatus* (Ylla et al., 2021) and *Laupala kohalensis* (Blankers et al., 2018; Ylla et al., 2021).

## 3 RESULTS AND DISCUSSION

### 3.1 Identification of sex-biased genes

From our assessment of expression across the 15,539 CDS in our main target species *G. bimaculatus* (Ylla et al., 2021), we report that sex-biased gene expression was most common in the gonadal tissues, where 4,822 (31.0%) of all *G. bimaculatus* genes under study were sex-biased in expression: 2,698 (17.4%) and 2,124 (13.7%) genes had ovary-biased and testis-biased expression respectively, and a total of 10,717 (69.0%) were unbiased in expression (Fig.2). By comparison, sex-biased gene expression was markedly less common in the somatic reproductive system, where only 5.6% of genes were sex-biased, with 353 (2.3%) and 520 (3.3%) genes showing female- and male-bias respectively. As compared to the gonad, markedly fewer genes exhibited female-biased and male-biased expression in the nervous system tissues, where 4.5% of 15,539 *G. bimaculatus* genes had sex-biased expression in the ventral nerve cord: 279 (1.8%) and 425 (2.7%) were female- and male-biased respectively (Fig. 2). For the brain, only 1.0% of genes were sex-biased in expression, with 51 (0.33%) and 106 (0.68%) being female- and male-biased respectively, an uncommonness that notably has also been suggested for brains of *D melanogaster* (Huylmans & Parsch, 2015). The patterns in *G. bimaculatus* were supported by strong correlations in FPKM among replicates with Spearman’s R≥0.92 across all studied genes (P<0.05; Fig. S1; one exception being the male somatic reproductive system, R=0.71, P<0.05), indicating high reproducibility of expression profiles.

Together, using the present criteria, it is evident that sex-biased gene expression is most common in the gonad, which is consistent with high phenotypic and transcriptional dimorphism of these sex organs in animals (Arbeitman et al., 2004; Parisi et al., 2004; Zhang et al., 2004; Small et al., 2009; Oliver et al., 2010; Meisel, 2011; Harrison et al., 2015; Whittle & Extavour, 2017; Whittle & Extavour, 2019; Whittle et al., 2020). In contrast, sex-biased gene expression is markedly less common in the somatic reproductive system and ventral nerve cord, and least common in the brain of *G. bimaculatus*.

### 3.2 Molecular evolution of sex-biased genes

#### 3.2.1 Rates of evolution

Following reciprocal BLASTX (Altschul et al., 1990) between *G. bimaculatus* and *G. assimilis* CDS and retention of genes with unsaturated dN and dS values (<1.5) after alignment, we identified 7,220 high confidence *G. bimaculatus*-*G. assimilis* orthologs that were used for all dN/dS analyses. Across all 7,220 orthologs under study, we found that the alignments with gaps removed were on average 68.0% (standard error=0.3%) of the original *G. bimaculatus* CDS length, and that the median dN/dS was 0.1152. The median dN was 0.0042 and median dS was 0.0396, values that were substantially <1, consistent with unsaturated substitution rates and a close phylogenetic relatedness between these two sister *Gryllus* species. Notably, the 90^th^ percentile of dN values was 0.042 and 95^th^ percentile was 0.094, also each well below 1, which facilitates precise ortholog detection (by a protein similarity search, reciprocal BLASTX (Altschul et al., 1997)), and indicates the studied ortholog gene set does not exclude relatively rapidly evolving genes in the genome(s) (that is, includes those with 22-fold higher dN than the median). Further, we found that the percent of all male-biased and female-biased *G. bimaculatus* genes (shown in Fig. 2) respectively that had high confidence orthologs between the two *Gryllus* species was 57.7% and 75.7% for the gonads, 55.2% and 52.1% for the somatic reproductive system, 42.4% and 39.2% for the brain, and 50.3% and 64.2% for the ventral nerve cord. Of note, the fact that we detected the fewest orthologs for the brain is suggestive of rapid protein sequence evolution of sex-biased genes in that tissue, which typically limits ortholog detection between divergent sequences (and/or sometimes may reflect gene losses/gains), while the highest detection in ovary-biased genes suggests putatively relatively slow protein sequence evolution. These ortholog datasets were subjected to dN/dS analyses as described below.

To precisely reveal the relationship between sex-biased gene expression for each individual tissue type and dN/dS, we identified genes that were sex-biased in expression in only one of the four female-male paired tissues (gonad, somatic reproductive system, brain or ventral nerve cord) and unbiased in all three remaining tissues in *G. bimaculatus*. These genes are hereafter denoted as tissue-specific sex-biased, or TSSB genes (N_TSSB_ values provided in Table S3). We emphasize that the TSSB status of a gene indicates that there is a tissue-specific sex difference in expression (has female-biased or male-biased status) that is not observed in other tissues (unbiased status in all other tissues), and does not imply that this gene is not expressed in any other tissue. Further, we identified those genes with universally unbiased expression in all four tissues types as a control (N=3,449; Table S3). The vast majority of the 7,220 genes (with orthologs in both species) fell into one of these two categories (94.5% had TSSB or universally unbiased status, while the remainder had mixed statuses among tissues).

#### 3.2.2 dN/dS of sex-biased genes in the four tissue types and pleiotropy

The dN/dS values of sex-biased_TSSB_ genes for each of the four paired *Gryllus* tissue types under study, and for universally unbiased genes, are shown in Fig. 3A. In turn, for completeness, the dN/dS values of all sex-biased genes for each tissue, regardless of status in other tissues (sex-biased_ALL_), are shown in Fig. 3B. The results show that in this cricket model, testis-biased_TSSB_ genes evolved faster than ovary-biased_TSSB_ and universally unbiased genes (Mann-Whitney U (MWU)-tests P<0.001 and 0.05 respectively, Fig. 3A). Further, sex-biased brain genes, while uncommon (Fig. 2, Table S3), evolved exceptionally rapidly. In particular, we noted faster evolution of the female-biased_ALL_ brain genes than of unbiased_ALL_ genes (MWU-test P=0.047, Fig. 3B). Given the P value is near the cutoff of 0.05, we analysed dN/dS of each brain gene set on a gene-by-gene basis for further scrutiny (see below section “*3.3 Rapid evolution of sex-biased genes from the brain*”). In turn, no statistically significant differences in dN/dS were observed among male-biased, female-biased, or unbiased genes from the somatic reproductive system or ventral nerve cords (using the TSSB genes and universally unbiased genes in Fig. 3A, or using ALL genes per tissue type in Fig. 3B (MWU-tests P>0.05)). In this regard, it is evident that the primary molecular evolutionary patterns in this cricket system include the rapid evolution of testis-biased genes and of sex-biased brain genes, particularly female-biased brain genes.

**Fig. 3.**
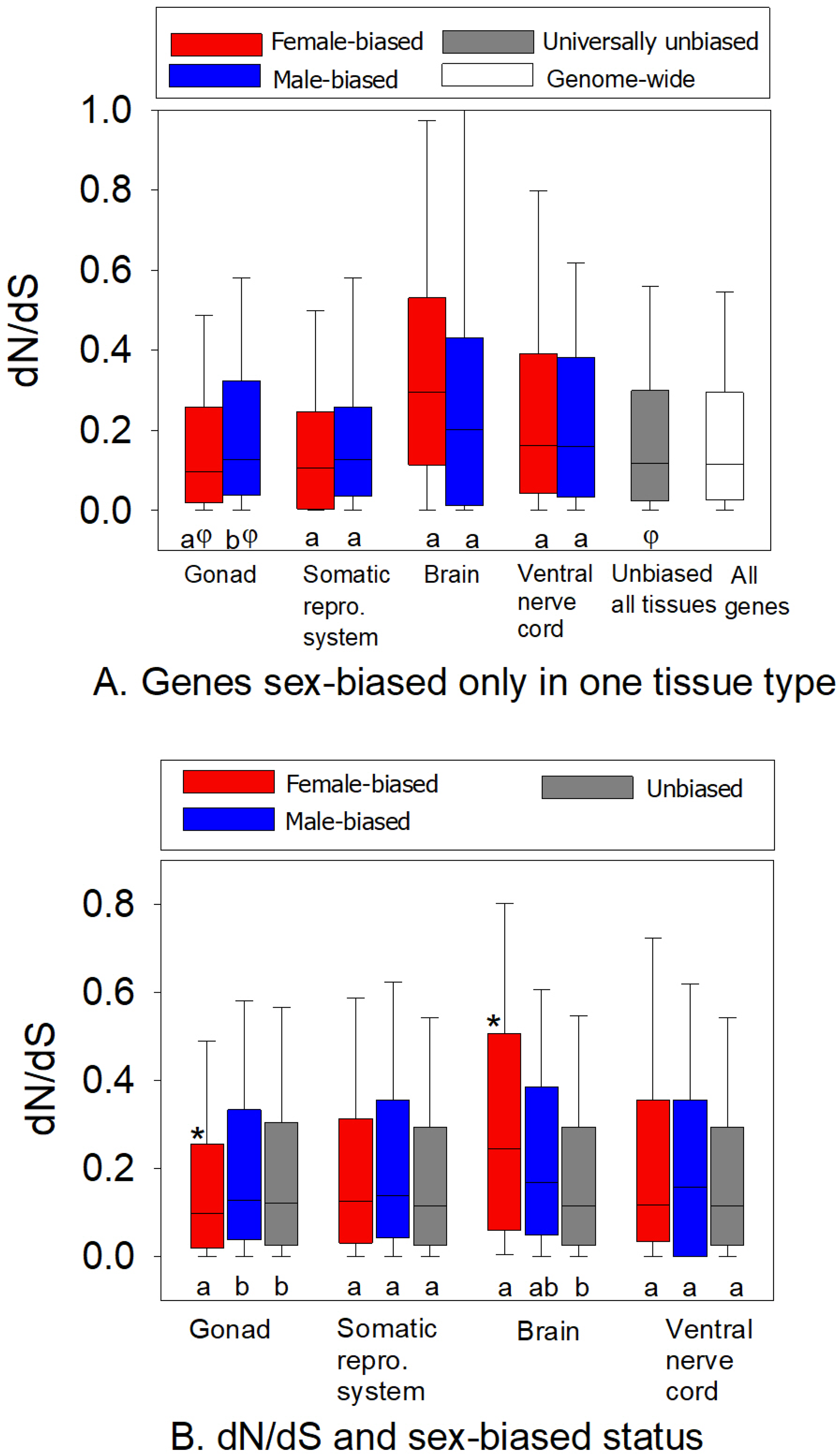
Box and whisker plots of the dN/dS values of genes with female- or male-biased expression in *G. bimaculatus*, and attained using the genes with high confidence orthologs its sister species *G. assimilus*. A) Genes with female- or male-biased gene expression in only one tissue type and unbiased in three remaining paired tissues, that is, with tissue-specific sex bias (TSSB). In addition, genes with universally unbiased expression in all four paired tissue types and the genome-wide dN/dS are shown. B) dN/dS of all (ALL) genes with sex-biased expression in each of four tissue types regardless of status in other tissues. In panel A, different letters (a, b) under the two bars within each tissue type indicate a statistically significant difference (MWU-test P<0.05), and φ indicates the difference in dN/dS in with respect to universally unbiased genes (MWU-tests P<0.05). For panel B, different letters among the three bars within each tissue type indicates MWU-test P<0.05 (note that “ab” for the brain indicates no difference of male-biased to female-biased or unbiased genes) and *indicates a difference in dN/dS between female-brain biased and ovary-biased genes. N values of genes per category are provided in Table S3. repro. = reproductive. Outliers above the 95^th^ percentile, including dN/dS >1, were excluded from the figure to allow visualizations on the Y-axis.

As a measure of pleiotropy, we examined the expression breadth across tissues (using nine tissues, the four paired female and male tissues and the male accessory glands, see Materials and Methods), which is thought to strengthen purifying selection and in turn may restrict adaptive evolutionary potential (Otto, 2004; Larracuente et al., 2008; Mank et al., 2008; Mank & Ellegren, 2009; Meisel, 2011; Assis et al., 2012; Dean & Mank, 2016; Whittle & Extavour, 2017). Genes were categorized into bins based on expression at >5 FPKM in 1-2, 3-4, 5-6, and 7-9 tissues. As shown in Fig. 4A, when studying all 7,220 genes with high confidence orthologs, we found that the rate of evolution of *Gryllus* genes was strongly inversely correlated with expression breadth. The lowest dN/dS values were found in genes transcribed in 7-9 tissues under study (median dN/dS=0.096), and the highest in genes expressed in 1-2 tissues (median=0.221, Ranked ANOVA and Dunn’s paired contrasts P<0.05). Further, as indicated in Fig. 4B, with respect to sex-biased gene expression, we found that testis-biased_TSSB_ genes had markedly lower expression breadth than ovary-biased_TSSB_ genes and than universally unbiased genes (MWU-tests P<0.001). Female-biased_TSSB_ brain genes had the smallest median expression breadth of all studied categories, which despite their low N value (Table S3), was statistically significantly lower than that of the universally unbiased genes (MWU-test P=0.021, Fig. 4B). Thus, this suggests a plausible connection between rapid protein sequence evolution and pleiotropy for sex-biased genes from the brain and the male gonad, either due to relaxed constraint in itself, and/or due to an associated freedom to evolve functional changes under low purifying constraint (see below section “*3.7 Evidence of A History of Positive Selection in Sex-Biased Gonadal and Brain Genes”*). In the following sections, we focus in detail on the dN/dS patterns for sex-biased genes in the brain and the reproductive system in Fig. 3 and Fig. 4, and consider further the putative roles of pleiotropy and positive selection in affecting their molecular evolution.

**Fig. 4.**
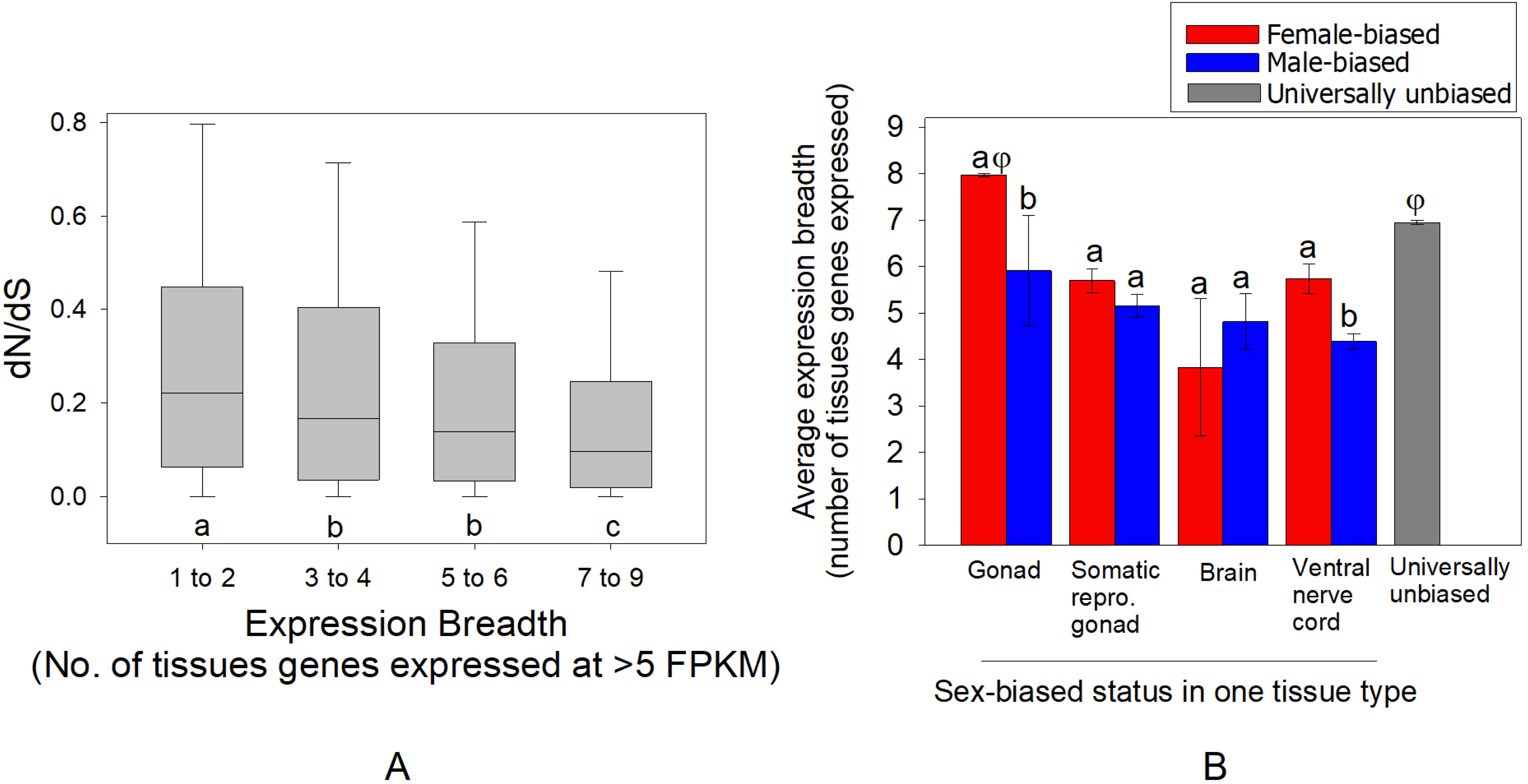
A) Box and whisker plots of the dN/dS values of all studied genes with respect to their expression breadth, or pleiotropy, in *G. bimaculatus* (N=7,220 genes). B) The average expression breadth (number of tissues with expression of a gene ≥5FPKM) of genes with sex-biased expression in only one tissue type, that is, with female- or male-biased_TSSB_ expression. In A, different letters below bars indicate a statistically significant difference using ranked ANOVA with Dunn’s paired contrast (P<0.05). In B, different letters in each pair of bars indicate a difference using MWU-tests. φ above ovary-biased and universally unbiased genes indicates a statistically significant difference from each other and from all other bars. Error bars in B indicate standard errors. repro. = reproductive. For panel A, outliers above the 95^th^ percentile, including dN/dS>1 were excluded for visualization on the Y-axis.

### 3.3 Rapid evolution of sex-biased genes from the brain

With respect to the brain, female-biased_TSSB_ genes had markedly higher median dN/dS values (median=0.295) than male-biased_TSSB_ genes (0.203, Fig. 3A), although that contrast was not statistically significant (MWU-test P=0.739). This may reflect the low statistical power of this comparison due to the rarity of genes with sex-biased_TSSB_ brain status (Table S3). When studying all genes with sex-biased_ALL_ expression in the brain, regardless of their expression status in other tissues (Fig. 3B), we found that the 20 female-biased_ALL_ brain genes had substantially higher median dN/dS values (median= 0.245) than the 45 male-biased_ALL_ (0.169) and the unbiased_ALL_ brain-expressed genes (0.115), wherein its contrast to the unbiased set was, as aforementioned, statistically significant (MWU-test P=0.047). Thus, the statistical tests suggest there are significant patterns in the brain (Fig. 3AB). Nonetheless, given the growing recognition that P-values alone may not always provide a full perspective to discern important biological patterns (Amrhein et al., 2019), particularly for samples with small sizes (such as for sex-biased brain genes studied here, Fig. 2), and given the close proximity of the core P value to 0.05, we aimed to further assess these findings by examining the sex-biased brain_ALL_ genes on a gene-by-gene basis, including their rates of evolution and their putative functions, as shown in Table 1. Using this approach, we show that 11 of the 20 female-biased_ALL_ brain genes (Fig. 3B) and 19 of 45 male-biased_ALL_ brain genes had dN/dS values more than two-fold higher (>0.236) than the median observed for universally unbiased genes (median=0.118; this value is shown in Fig. 3A; median across the whole genome=0.115). This close examination of individual genes within each gene set, combined with the observed P-values (Fig. 3), taken together indicate that the sex-biased brain genes share a striking propensity to evolve rapidly as compared to universally unbiased genes and the genome as a whole, with the effect being particularly elevated in the female brain (Fig. 3A, Table 1). While the study of protein evolution of sex-biased brain genes (brains *sensu stricto*, rather than simply heads, or pooled brain-eye tissues as considered by some previous studies (Catalan et al., 2018; Congrains et al., 2018)) remains rare, rapid evolution of female-biased brain genes has been reported in some bird embryos (Mank et al., 2007), and in some autosomal genes in flies (Khodursky et al., 2020). However, an opposite pattern of rapid evolution of male-biased brain genes for several stages of development was reported in humans (Shi et al., 2016). The avian result was interpreted as possibly reflecting selective pressures arising from brain-regulated mating behaviors (Mank et al., 2007). We suggest that this may also be a significant factor contributing to the trend of rapid evolution of sex-biased brain genes here for crickets.

**Table 1.**
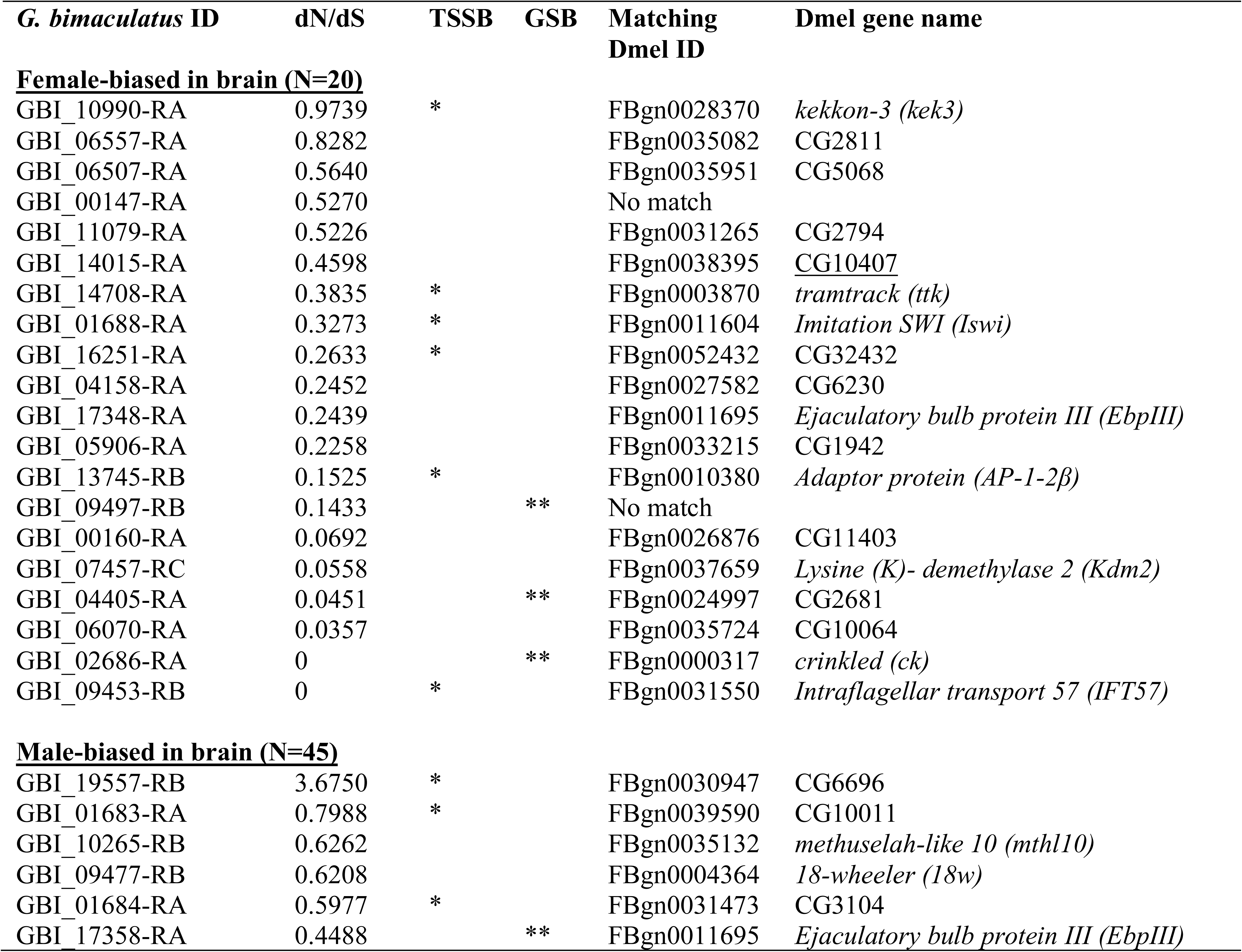
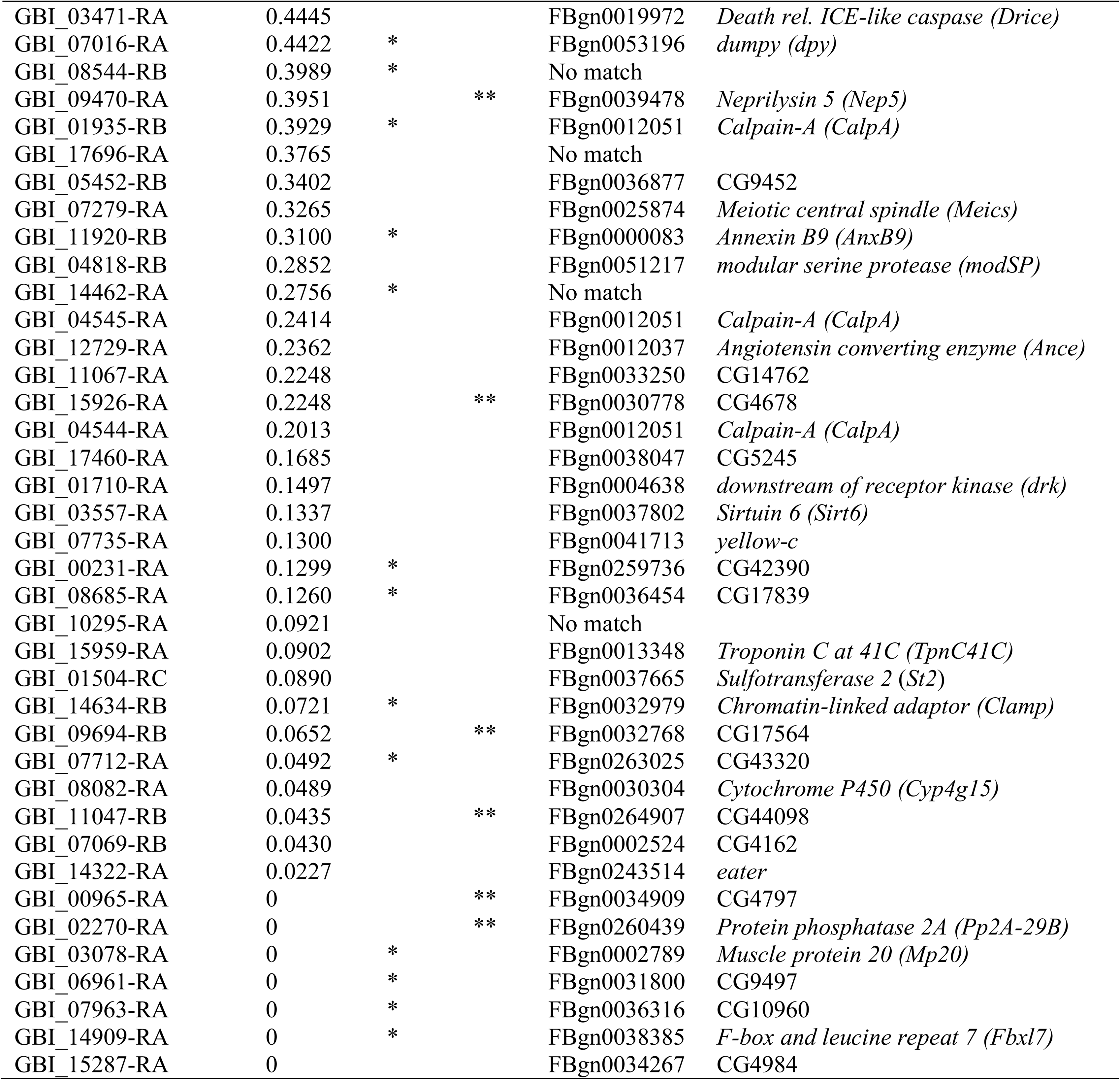
The dN/dS values of all female-biased brain genes and male-biased brain genes among the 7,220 genes with *G. bimaculatus* and *G. assimilis* orthologs. Tissue-specific sex bias (TSSB) indicates genes that have sex-biased expression in the brain and are unbiased in all other paired tissues (shown by “*”). Gonad sex bias (GSB) indicates the gene has the same female- or male-biased expression status in the gonad as in the brain and is unbiased in other tissues (“**”). The best matching *D. melanogaster* (Dmel) ortholog is shown with identifiers and gene names from FlyBase (Gramates et al., 2017). Genes are listed by highest to lowest dN/dS values per category.

We examined the putative GO functions for the sex-biased brain genes (Fig. 3). For this, we used single-direction BLASTX (Altschul et al., 1990) of the *G. bimaculatus* entire CDS list to the CDS of well-studied insect model *D. melanogaster* (Gramates et al., 2017) to identify its putative orthologs, which were assessed in the GO tool DAVID (Huang da et al., 2009) (note that single direction BLASTX was used for functional analysis, rather than the reciprocal BLASTX approach that was used for *G. bimaculatus* and *G. assimilis* contrasts for dN/dS, see details in Materials and Methods). First, we conducted enrichment analyses using all *G. bimaculatus sex*-biased brain genes, regardless of the two-species *Gryllus* ortholog status (N values in Fig. 2). We found that female-biased brain genes were enriched for transcriptional functions and sensory perception, while male-biased brain genes were enriched for proteolysis and neuron remodelling (Table S4). We then identified putative functions of those genes with orthologs that were used in our dN/dS analyses, including ALL (sex-biased) genes and the subset of genes that had TSSB status (Fig. 3AB, Table S3) on a gene-by-gene basis as shown in Table 1 (note: any brain genes that had the same sex-biased expression status in the gonad are also shown as gonad sex bias “GSB”). We observed that the predicted functions of female-biased brain genes included involvement in neurotransmission (*AP-1-2β*), apoptosis (*D. melanogaster* ID number CG2681), and DNA binding (CG11403) (Table 1). Remarkably, certain brain-expressed genes were predicted to be involved in sexual processes or organs, including multicellular reproduction (CG10407), inter-male aggressive behavior (*tramtrack*) (Yamamoto et al., 1998) and the ejaculatory bulb (*EbpIII*) (Table 1). These genes had exceptionally elevated dN/dS values of 0.460, 0.384 and 0.244 respectively (Table 1), as compared to the median for universally unbiased genes (median=0.118, Fig. 3A). The fastest evolving female-biased brain gene (dN/dS=0.970) was a putative ortholog of *kekkon-3*, a member of a *kekkon* gene family known to be involved in neuron function and differentiation of the central nervous system in flies (Musacchio & Perrimon, 1996), that is conserved in flies and mosquitoes (MacLaren et al., 2004). Collectively, the genes that are upregulated in the cricket female brain may play significant roles in female behaviors, such as mating functions, possibly contributing to their rapid divergence.

Despite a tendency for accelerated evolution, not every female-biased *G. bimaculatus* brain gene evolved rapidly (Table 1). For instance, one highly constrained gene (GBI_02686-RA, (dN/dS=0 (dN=0 dS=0.041)) was an ortholog match to *D. melanogaster crinkled*, which is involved in hearing (vibration sensing) in both flies and vertebrates (Todi et al., 2005; Boekhoff-Falk & Eberl, 2014). We speculate that a history of strong constraint reflected in dN/dS of this female-biased brain gene could indicate an essential role of negative phonotaxis (potentially relevant to avoiding predators (Schneider et al., 2017)), perhaps an effect enhanced in females. However, the sex-biased expression of this putative *crinkled* gene may also suggest it has a sexual role. A fundamental factor underlying male-female attraction in *G. bimaculatus* is song, which is used by males to attract females (positive phonotaxis), and is thought to be regulated by the auditory neural pathways involving the brain (Lankheet et al., 2017; Sakai et al., 2017). Thus, it is tempting to speculate that the strong purifying selection on this particular female-biased gene could partly reflect an essential role in receiving male auditory signals for reproduction, courtship and mating. Further studies in crickets should assess sex-biased gene expression in the brain of males and females from mixed mating populations (virgin males and females were studied herein, see Materials and Methods) to identify brain-related auditory genes potentially involved in mating. Questions of interest for future work include whether these genes tend to be highly conserved in sequence, and/or whether some may exhibit adaptive changes possibly due to neural-related mating behaviors. Studies in related crickets (*Teleogryllus*) have suggested that neural genes involved in mating, including those involved in acoustics, may have key roles in early stages of male or female development (Kasumovic et al., 2016), and be associated with sex-related behavioral plasticity and abrupt adaptive evolutionary changes (Pascoal et al., 2020). Thus, acoustics, mating, and neural gene sequence evolution may be intrinsically tied. Additional valuable future directions could include study of sex-biased expression in the male and female auditory organs located on the tibia of the forelegs in crickets (Lankheet et al., 2017; Schneider et al., 2017), in the antennae, which are involved in male-female attraction and male-male aggression and contain neurons involved in sex-related pheromonal signalling (Murakami & Itoh, 2003; Yoritsune & Aonuma, 2012; Boekhoff-Falk & Eberl, 2014), and in the terminal abdominal ganglion, which has been linked to mating behaviors (Sakai et al., 2017). These types of follow-up studies in *G. bimaculatus* will help further identify and evaluate the evolutionary roles of brain and neural genes linked to mating and sex-related auditory and pheromonal signalling in this taxon.

With regard to the male-biased brain genes, a range of predicted functions were observed. For instance, multiple genes were associated with phagocytosis (six of 45 genes), and early-stage development (three genes). In addition, some genes had predicted sexual roles. In particular, a putative *G. bimaculatus* ortholog (GBI_17358-RA) of a *D. melanogaster* ejaculatory bulb protein *EbpIII* had a dN/dS value of 0.449, which was nearly four-fold higher than the median for universally unbiased genes (0.118, Table 1). This same *EbpIII* related gene (GBI_17358-RA) was also found to be testis-biased in expression (Table 1), which is consistent with putatively significant roles in both brain and testicular functions in *G. bimaculatus*. As described above, a different *G. bimaculatus* gene (GBI_17348-RA) that was also an ortholog match to *D. melanogaster EbpIII* was sex-biased in the female-brain (dN/dS=0.243, Table 1), suggesting the possibility that there are two distinct paralogs to this gene, which may have different roles in male and female brains in crickets. These two genes matching *EbpIII*, one biased in the male-brain and the other in the female brain, are candidates to be involved in male-female attraction, mating or sexual behaviors. In *D. melanogaster,* while the exact functions of *EbpIII* remain under assessment, its key predictive classifications include olfactory function, post-mating behavior, and mating plugs (flybase.org, (Gramates et al., 2017)), further suggesting a possible function in male-female brain mediated sexual behaviors in *G bimaculatus*. We also discovered that the male-biased brain genes included a putative ortholog of *Angiotensin converting enzyme*, a gene whose functions include involvement in *D. melanogaster* spermatid nucleus differentiation and sperm individualization (Hurst et al., 2003). This gene had a dN/dS value of 0.236, which is double the median of universally unbiased genes (Table 1). In this regard, multiple male-biased brain genes exhibit rapid protein-level divergence and are candidates to have potential sex-related roles in this taxon.

While the tendency for rapid protein sequence evolution of sex-biased brain genes in Table 1 could largely result from relaxed purifying constraint and neutral protein sequence changes, as suggested by their low pleiotropy (Fig. 4B), the low pleiotropy could in principle also act to accelerate protein changes by more readily allowing adaptive functional changes (Otto, 2004; Larracuente et al., 2008; Mank et al., 2008; Mank & Ellegren, 2009; Meisel, 2011; Assis et al., 2012; Dean & Mank, 2016; Whittle & Extavour, 2017). We suggest here that several features of the mating biology of *G. bimaculatus* might cause episodic adaptive evolution and underlie the high dN/dS values observed herein (see also below section “*3.7 Evidence of A History of Positive Selection in Sex-Biased Gonadal and Brain Genes*”). For instance, *G. bimaculatus* exhibits aggressive male-male fighting and mate guarding (Vedenina & Shestakov, 2018; Gee, 2019) and males transfer larger spermatophores to females when in the company of rival males (Lyons & Barnard, 2006). Such behaviors are likely mediated by the male brain. This could, in principle, lead to sexual selection pressures on the male-biased brain genes, which might give rise to adaptive changes in dN/dS. It is also feasible that inter-locus sexual conflict could contribute to the tendency for rapid evolution of both sets of male- and female-biased brain genes (Koene et al., 2013; Mank et al., 2013; Pennell et al., 2016). In other words, it is possible that aggressive male-male behaviors in *G. bimaculatus* (Vedenina & Shestakov, 2018; Gee, 2019), directed by male-biased brain genes, may negatively affect female fitness. This might be predicted to lead to an adaptive response in female-biased brain genes (e.g., genes regulating the behavior of removal of spermatophores of certain males by females after mating (Bateman et al., 2001)), causing an evolutionary “arms race” that could in theory accelerate evolution of proteins of both types of genes (Ellegren & Parsch, 2007; Mank et al., 2013). Taken together, we suggest that there are several plausible mechanisms related to mating biology of this taxon that may underlie the observed patterns for sex-biased brain genes (Table 1), mediated by low pleiotropy and, in turn, an enhanced potential for adaptive evolution.

A key aspect of future research should include studies of male and female brains in courtship and mating environments, given that the brain likely regulates these sex-related behaviors in *Gryllus* including song, sexual attraction, copulation and aggression (Matsumoto & Sakai, 2000; Haberkern & Hedwig, 2016; Sakai et al., 2017), and that brain expression has been found to differ between sexes under mating conditions in other insects such as *Drosophila* (based on expression analysis of combined whole head-thorax expression in males and females in that study (Fowler et al., 2019)). We anticipate that in crickets under courtship and mating environments, more genes, in addition to those identified in for virgins (Table 1), may exhibit sex-biased expression given the intense male competition (Vedenina & Shestakov, 2018; Gee, 2019) and the propensity for female-choice in this taxon (Bateman et al., 2001; Zhemchuzhnikov et al., 2017). In turn, future research in these crickets may allow further testing of the notion that mating behaviors may underlie the rapid protein sequence evolution of some brain genes, and thus ultimately possibly contribute to processes such as reproductive isolation and speciation.

It should be recognized that while sex-biased brain genes, by definition, exhibit differences in gene expression between the female and male brain, these sex biases may reflect differences in cellular expression and/or allometric scaling differences in male and female brains. As an example, the female-biased brain gene *crinkled* (Table 1) may be more highly expressed in all female than male brain cells, the female brain may typically contain more cells that express this gene (Montgomery & Mank, 2016), and/or the gene may be more highly expressed in cells from a particular sub-region(s) of the brain (Tuller et al., 2008), whereby the size or cell composition of the subsections may vary between females and males (Montgomery & Mank, 2016). Further studies of gene expression, and the allometry of subsections of the brain in males and females, would be needed to distinguish among these possibilities, and to better understand the factors underlying differences in male and female brain expression.

### 3.4 Rates of Evolution of Sex-biased Genes from the Reproductive System

#### 3.4.1 Rapid evolution of testis-biased genes

With respect to sex-biased expression in the gonads and dN/dS, which has been more commonly studied as compared to the brain in insects, we observed marked differences in rates of protein sequence evolution among sex-biased_TSSB_ genes. First, dN/dS decreased progressively from testis-biased_TSSB_ (median=0.128), to universally unbiased genes (median=0.118) to ovary-biased genes (median=0.097, each paired MWU-test P<0.05; see also Fig. 3B). Thus, the rate differences were most marked between testis-biased_TSSB_ and ovary-biased_TSSB_ genes, with intermediate values for those with universally unbiased expression. The tendency for rapid evolution of testis-biased genes in this cricket concurs with patterns observed for *Drosophila* (Zhang et al., 2004; Proschel et al., 2006; Ellegren & Parsch, 2007; Zhang et al., 2007; Jiang & Machado, 2009; Meisel, 2011; Assis et al., 2012; Perry et al., 2015; Grath & Parsch, 2016; Whittle & Extavour, 2019) (see results in a related fly (Congrains et al., 2018)), and recent findings in beetles (*Tribolium castaneum*) (Whittle et al., 2020). However, the results are opposite to the rapid evolution of ovary-biased (or ovary-specific) genes previously reported in the mosquitoes *Aedes* and *Anopheles* (Papa et al., 2017; Whittle & Extavour, 2017). In this regard, it is worth considering possible reasons for variation in the effects of sex-biased gonadal expression among these insect taxa.

Given that *Gryllus* (Orthoptera) is a distant outgroup to the two Diptera groups (*Drosophila* and *Aedes/Anopheles*) and the Coleoptera (*Tribolium*) (Misof et al., 2014) it may be suggested, based on the collective anecdotal evidence, that there could be a shared ancestral effect of testis-biased expression in *Drosophila-Tribolium-Gryllus* (Zhang et al., 2004; Ellegren & Parsch, 2007; Harrison et al., 2015; Whittle et al., 2020)) and a derived effect of rapid evolution of ovary-biased (or ovary-specific) genes in *Aedes/Anopheles* (Papa et al., 2017; Whittle & Extavour, 2017). Under this hypothesis, the pattern observed for studied *Aedes* and *Anopheles* species would be a derived feature, and could reflect variation in mating biology among these insects. For example, although both *Drosophila* and *Aedes aegypti* (the *Aedes* species studied in (Whittle & Extavour, 2017)) are polyandrous and thus prone to sperm competition, the polyandry is thought to be relatively weak in the mosquitoes (Helinski et al., 2012). Further, this mosquito can exhibit intensive male swarming during courtship that may involve female-female mosquito competition and/or male-mate choice (Oliva et al., 2014; Whittle & Extavour, 2017). In addition, nonporous mating plugs are formed in the female mosquito reproductive tract after mating, which prevent sperm competition (Oliva et al., 2014) and thus differ both from the mating plugs formed in *Drosophila*, which allows sperm transfer from competitor males (Manier et al., 2010; Avila et al., 2015), and from observations of complete sperm mixing from multiple males in *Gryllus* (Simmons, 1986). Any of these mating-related features could in principle give rise to sexual selection and the relatively faster evolution of ovary-biased than testis-biased genes in mosquitoes (Whittle & Extavour, 2017), and not in the other studied insects. In addition, relaxed purifying selection, possibly due to low pleiotropy, may be more common for ovary-biased genes in the mosquitoes (Whittle & Extavour, 2017), as inferred for testis-biased (or male-biased) genes in some organisms, including flies (Allen et al., 2018; Ghiselli et al., 2018), and suggested for the crickets studied here (Fig. 4B). Studies in even more insect models, particularly in monogamous versus polyandrous species (Harrison et al., 2015), and in additional insects with various degrees of male-male or female-female competition and with and without impermeable mating plugs (Whittle & Extavour, 2017), would help elucidate whether and how and why the effects of sex-biased transcription on protein evolution vary among insects.

Functional predictions of testis-biased_TSSB_ and ovary-biased_TSSB_ genes in *G. bimaculatus* are shown in Table 2 (using *D. melanogaster* orthologs and GO clustering). Testis-biased_TSSB_ genes were predicted to be preferentially involved in cilium functions, potentially reflecting roles in sperm motility (Trotschel et al., 2019). Ovary-biased_TSSB_ genes were particularly involved in fundamental processes such as transcription functions. Thus, the former may be linked to specialized functions of the male gonad, and sperm functionality, while the latter may include genes involved in broader functions in addition to their roles in the female gonad. In terms of GO functions of the universally unbiased genes, these genes were preferentially involved in core cellular and nuclear functions including protein structure (coiled coil), nucleotide binding and splicing (Table S5), differing from more specialized functions of testis-biased genes.

**Table 2.** Top GO functional groups for testis-biased_TSSB_ and ovary-biased_TSSB_ genes identified in *G. bimaculatus* (those with orthologs in *G. assimilis*). Genes were sex-biased only in the gonads and not in the somatic reproductive system, brain or ventral nerve cords (tissue-specific sex-biased, TSSB). The top clusters with the greatest enrichment (abundance) scores are shown per category. *P*-values are derived from a modified Fisher’s test, where lower values indicate greater enrichment. Data are from DAVID software (Huang da et al., 2009) using those *G. bimaculatus* genes with predicted *D. melanogaster* orthologs.

| **Ovary-biased genes (N=1,858)** |  | **Testis-biased genes (N=1,055)** |  |
| --- | --- | --- | --- |
| **GO Function** | **P-value** | **GO Function** | **P-value** |
| **Cluster 1: Enrichment Score 10.31** |  | **Cluster 1: Enrichment Score: 5.38** |  |
| nucleotide-binding | 1.00E-15 | ubiquitin-protein transferase activity | 1.20E-07 |
| ATP-binding | 2.00E-14 | **Cluster 2: Enrichment Score: 3.66** |  |
| **Cluster 2: Enrichment Score 7.19** |  | cilium assembly | 3.70E-06 |
| WD40/YVTN repeat-like-containing domain | 7.70E-09 | cilium morphogenesis | 6.90E-05 |
| **Cluster 3: Enrichment Score 5.41** |  | **Cluster 3: Enrichment Score: 3.28** |  |
| transcription, DNA-templated | 5.70E-03 | Nucleotide-binding | 2.50E-04 |
|  |  | ATP binding | 6.40E-04 |

It is worth mentioning that in Fig. 3A, while testis-biased_TSSB_ genes had higher dN/dS values than ovary-biased_TSSB_ genes and than the universally unbiased genes, they did not exhibit any statistically significant differences with respect to the male-biased genes from the three other tissues, including from the brain (MWU-tests P>0.05). Significantly, however, given the much greater abundance of testis-biased_TSSB_ genes than male-biased_TSSB_ genes from other tissues (8- to 65-fold more common, Fig. 2, Table S3), it may be inferred that testis-biased gene expression plays a substantial role in shaping the portion of the genome that is rapidly evolving in *G. bimaculatus*.

#### 3.4.2 Sex-biased gonadal expression in G. assimilis

While our main target for expression analyses was *G. bimaculatus*, and *G. assimilis* was used primarily as a reference point to measure rates of protein divergence, we considered the degree of conservation of gene expression between the two species for the 7,220 genes with orthologs for the gonads (which had the largest N values of all tissues, Table S3). The results are shown in Fig. S2 and are described in Text File S1. We observed that the finding of elevated dN/dS of testis-biased versus ovary-biased genes was robust to whether the sex-biased status (testis-biased, ovary-biased) was observed in one species or was conserved in both of these species. Thus, testis-biased expression in one species (i.e., *G. bimaculatus* or *G. assimilis*, Fig. S2) is sufficient to predict elevated pairwise dN/dS.

#### 3.4.3 Possible influence of the faster-X effect

The faster-X theory contends that genes located on the X-chromosome evolve faster than those on autosomes in male heterogametic XY systems due to rapid fixation of recessive beneficial mutations in hemizygous males (or the Z-chromosome in WZ systems) (Charlesworth et al., 1987). A faster-X effect could also possibly result from relaxed selection on the X-chromosome as compared to autosomes due to lower effective population size (Parsch & Ellegren, 2013). The former cause of a faster-X effect may be evidenced by rapid evolution of male-biased (or typically testis-biased) genes as compared to female-biased and unbiased genes, while the absence of this relationship among sex-biased genes may suggest relaxed selection (Mank et al., 2010a; Parsch & Ellegren, 2013). Given that the recently available and large (1.66 Gbp) *G. bimaculatus* genome remains on scaffolds in this non-traditional model (Ylla et al., 2021), that hypothesis cannot yet be explicitly tested, unlike in insect taxa with widely available and intensively studied genomes (e.g. *Drosophila, Tribolium* (Mank et al., 2010b; Whittle et al., 2020)). Nonetheless, it is worthwhile to consider whether the faster-X effect could contribute to any of the results herein. A recent study of the faster-X effect in beetles (*Tribolium*, an X/Y system) found weak or absent male dosage compensation in the gonads of that taxon, which was associated with an excess of female-biased gonadal genes on the X-chromosome, and the X-chromosome exhibited lower dN/dS than the autosomes (Whittle et al., 2020). These observations suggested an absence of a faster-X effect in *Tribolium*, possibly mediated by low gonadal dosage compensation and rarity of X-linked male-biased genes. A weak faster-X effect has been suggested in *Drosophila* (Mank et al., 2010b; Meisel & Connallon, 2013; Avila et al., 2014; Charlesworth et al., 2018), possibly due to poor dosage compensation in gonads of that taxon (Gu & Walters, 2017; Argyridou & Parsch, 2018). In the XX (female) and X0 (male) system of aphids, a faster-X effect was observed, believed to arise under the selective non-neutral model (Jaquiery et al., 2018), and thus presumably male dosage compensation. Thus, this faster-X pattern could in principle also occur in the XX and X0 system of *G. bimaculatus* (Yoshimura et al., 2006). In this context, given that studied crickets and locusts (Camacho et al., 2015; Pascoal et al., 2020) including *G. bimaculatus* (Yoshimura et al., 2006) have cytologically relatively large X-chromosomes compared to the autosomes, we suggest that under specific circumstances, a faster-X effect could possibly give rise to the rapid evolution of testis-biased genes (as compared to ovary-biased and universally unbiased) found herein. Specifically, if there is full gonadal dosage compensation (or overcompensation) on the X chromosome in males in this cricket species then that may cause a high concentration of male-biased gonadal genes on the X chromosome. If there are few testis-biased genes on autosomes, then a faster-X effect could contribute at least partly to the observed patterns of highest dN/dS in testis-biased genes, with lower values for ovary-biased and unbiased genes (Fig. 3), a pattern expected under a selection-based faster-X effect (Parsch & Ellegren, 2013). Importantly, however, as here we have sex-biased expression data from the brain, we also suggest from our findings (Fig. 3, Table 1) that if brain genes are preferentially linked to the X chromosome and exhibit full dosage compensation, this could contribute to rapid evolution of male-biased brain genes (relative to unbiased genes), but could not give rise to the rapid evolution of female-biased brain genes, given that those genes are not monosomic (not X0) in females, excluding a putative role of a faster-X effect. Further studies will thus be valuable to deciphering whether the faster-X effect, and gonadal and brain dosage compensation, may contribute in some manner towards the observed rapid evolution of the testis-biased genes and male-biased brain genes in the cricket model.

### 3.5 Sex-biased genes from the somatic reproductive system

In contrast to the gonad, the lack of differences in dN/dS of male-biased_TSSB_ and female-biased_TSSB_ genes, and between those groups and the universally unbiased genes, for the somatic reproductive system (MWU-tests P>0.05, Fig. 3A; and when using ALL genes, Fig. 3B) is surprising, given the roles of these sexual tissues in reproductive success and fitness, including for the female tissues (oviducts, spermathecae, and bursa). Few comparable insect data of sex-biases in somatic reproductive system tissues are available. Some specific genes involved in the female reproductive tract in *Drosophila* have been linked to rapid and/or adaptive evolution, which may be due to their dynamic roles in receiving and maintaining sperm after mating (Swanson & Vacquier, 2002; Swanson et al., 2004) (note: see section “*3.7 Evidence of A History of Positive Selection in Sex-Biased Gonadal and Brain Genes*” which suggests a small number of female somatic reproductive system genes evolve adaptively). However, a separate assessment of genes broadly defined as female reproductive tract proteins in *D. melanogaster* (based on expression data from mixed or mated flies) showed those genes exhibited slow protein evolution (dN/dS), below the genome-wide average (Haerty et al., 2007). Our results from unmated *Gryllus* suggest no consistent differences in dN/dS between female-biased_TSSB_ somatic reproductive system genes and the universally unbiased genes or the genome as a whole (Fig. 3).

It is also notable that markedly fewer genes were sex-biased in expression in the somatic reproductive system as compared to the gonads (Fig. 2). One possible reason is that there may be an inherent variation in expression among individuals for the male somatic reproductive system (which had the least strongly correlated FPKM among replicates of all nine tissue types, Fig S1H), such that a consistent male to female difference in expression may be less apt to be observed for those tissues. Another possibility is that the gonads in adults are continuously supporting the dynamic process of gametogenesis (Pauli & Mahowald, 1990; Williamson & Lehmann, 1996) causing high female and male expression differentiation (Fig. 2), while the somatic reproductive system, particularly in unmated tissues as studied here, may be less dynamic, and thus exhibit less potential for differential transcription between males and females.

### 3.6 Rapid divergence of genes from the male accessory glands and seminal fluid proteins

For thoroughness in the study of reproductive structures, given that genes from the male accessory glands, including seminal fluid protein (SFPs), have been linked to rapid evolution in species of *Drosophila* (Haerty et al., 2007; Sepil et al., 2019), and in some identified cricket SFPs based on partial gene sets attained from assembled reproductive transcriptome sequences for species such as *G. firmus*, *G. pennsylvanicus* and *Allonemobius fasciatus* (Andres et al., 2006; Braswell et al., 2006; Andres et al., 2013), we assessed expression and evolution of such genes in *G. bimaculatus*. The findings for the male accessory glands (described in detail in Text File S1 and Table S6) showed that *G. bimaculatus* genes that had expression solely in the male accessory glands rarely had a high confidence ortholog in its sister species *G. assimilis*. Thus, this suggests a history of rapid evolution potentially so extensive that it prevents protein similarity detection by these methods, and/or a history of lineage-specific gene losses or gains of genes involved in this particular sexual tissue (Haerty et al., 2007; Tautz & Domazet-Loso, 2011).

For the study of SFPs, we used the recently available gene list of 134 SFPs from the species *D. melanogaster* as the reference (Sepil et al., 2019). The results are described in Text File S1 and Table S7. We found that only 20 *D. melanogaster* SFP genes had identifiable putative orthologs in *G. bimaculatus* (14.9%). Seven of those were included among the subset of 7,220 genes with between-species orthologs in the two species of *Gryllus* (note the stringent criteria used for the intra-*Gryllus* ortholog matches, see Materials and Methods). The dN/dS values of these seven genes are shown in Table 3; all were above the genome-wide median dN/dS value (0.115). Positive selection was indicated for the gene matching an odorant binding SFP protein *Obp56g*, with dN/dS>1 (Table 3). Together, we conclude that the putative SFPs in the crickets studied herein have evolved very rapidly, a feature shared with SFPs of *D. melanogaster* (Haerty et al., 2007; Sepil et al., 2019), and that could be due to their potential subjection to sex-related selection pressures. For instance, in flies SFPs may enhance sperm competitive ability in the female reproductive tract or egg release from the ovary (Heifetz et al., 2000; Fedorka et al., 2011), and males may alter relative production of different SFPs when exposed to male rivals (Fedorka et al., 2011). If similar types of mechanisms of sexual selection exist in crickets, then they could contribute to fast evolution of SFP genes. Another potentially significant behavioural factor in *G. bimaculatus*, is the tendency of females to preferentially retain deposited spermatophores of certain (larger) males (Simmons, 1986; Bateman et al., 2001), which comprises a mechanism of female-choice in this species (Bateman et al., 2001), potentially accelerating SFP evolution.

**Table 3.**
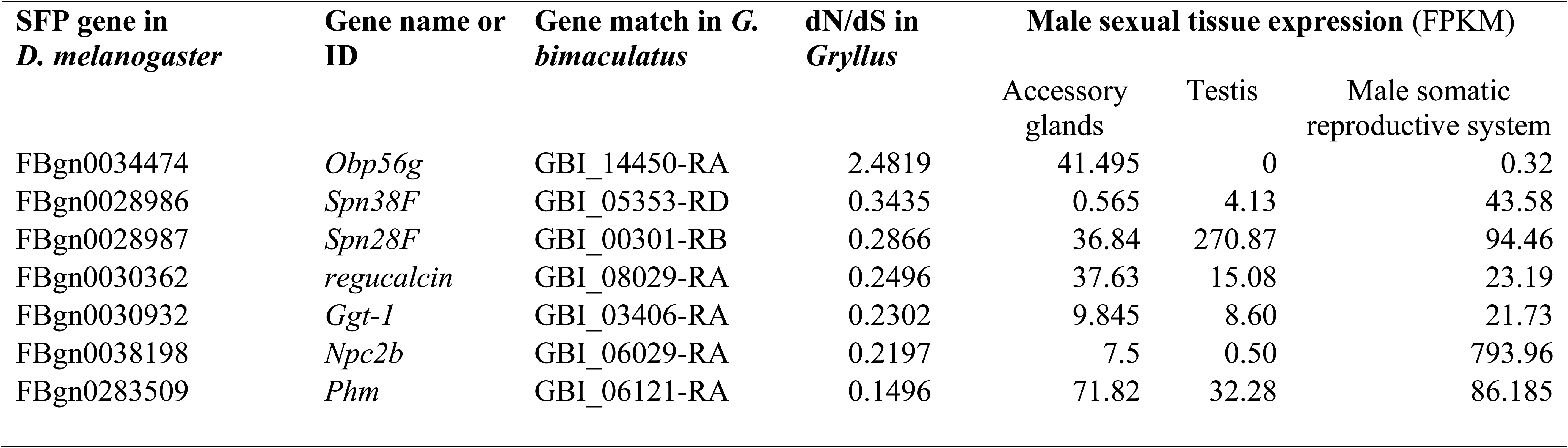
The *D. melanogaster* seminal fluid proteins (SFPs) (Sepil et al., 2019) that were found to have putative orthologs in *G. bimaculatus* (GB) among the subset of 7,220 genes with intra-*Gryllus* orthologs used for dN/dS analysis. Expression levels (FPKM) for each gene are shown for the three male sexual tissues under study.

| SFP gene in<br>*D. melanogaster* | Gene name or<br>ID | Gene match in *G.*<br>*bimaculatus* | dN/dS in<br>*Gryllus* | Male sexual tissue expression (FPKM) |  |  |
| --- | --- | --- | --- | --- | --- | --- |
|  |  |  |  | Accessory<br>glands | Testis | Male somatic<br>reproductive system |
| FBgn0034474 | *Obp56g* | GBI_14450-RA | 2.4819 | 41.495 | 0 | 0.32 |
| FBgn0028986 | *Spn38F* | GBI_05353-RD | 0.3435 | 0.565 | 4.13 | 43.58 |
| FBgn0028987 | *Spn28F* | GBI_00301-RB | 0.2866 | 36.84 | 270.87 | 94.46 |
| FBgn0030362 | *regucalcin* | GBI_08029-RA | 0.2496 | 37.63 | 15.08 | 23.19 |
| FBgn0030932 | *Ggt-1* | GBI_03406-RA | 0.2302 | 9.845 | 8.60 | 21.73 |
| FBgn0038198 | *Npc2b* | GBI_06029-RA | 0.2197 | 7.5 | 0.50 | 793.96 |
| FBgn0283509 | *Phm* | GBI_06121-RA | 0.1496 | 71.82 | 32.28 | 86.185 |

### 3.7 Evidence of A History of Positive Selection in Sex-Biased Gonadal and Brain Genes

Finally, we considered the incidences of positive selection among the 7,220 genes with between-species *Gryllus* orthologs. Gene-wide dN/dS>1 was taken as evidence of positive selection (Swanson et al., 2001; Torgerson et al., 2002; Nielsen et al., 2005; Clark et al., 2006; Yang, 2007; Hunt et al., 2011; Buschiazzo et al., 2012; Ghiselli et al., 2018; Hill et al., 2019)). The use of dN/dS>1 across a gene is a conservative means to identify positive selection (Swanson et al., 2001; Buschiazzo et al., 2012), as nonsynonymous codon changes should be sufficiently common to cause the ratio to exceed 1. We found that 1.63% of all the 7,220 *G. bimaculatus-G. assimilis* gene orthologs (N=118 genes) showed dN/dS>1.

We then considered whether dN/dS values of the sex-biased_TSSB_ genes from the gonad (Table 4), which had the highest N values of all tissues analysed (Table S3), were consistent with the aforementioned hypothesis that reduced gene pleiotropy, or expression breadth (and thus purifying selection), may lead to an enhanced opportunity for functional evolution of genes (Otto, 2004; Larracuente et al., 2008; Mank et al., 2008; Mank & Ellegren, 2009; Meisel, 2011; Assis et al., 2012; Whittle et al., 2020). We found that the percent of genes with positive selection increased from ovary-biased_TSSB_ genes (1.02%, 19 of 1,858) to universally unbiased genes (1.91%, 66 of 3,449) and testis-biased_TSSB_ genes (2.09%, 22 of 1,055; Chi^2^ P with Yates’ correction was <0.05 for each paired contrast to ovary-biased_TSSB_ genes, Table 4). In turn, expression breadth of these genes decreased from all ovary-biased_TSSB_ (average expression breadth of 7.97±0.04 (standard error)), to universally unbiased (6.95±0.05) and to testis-biased_TSSB_ genes (5.90±0.18 tissues; (MWU-tests P<0.001 for each of three paired contrasts (Fig. 4B). Strikingly, the differences were even more magnified in the subset of genes with dN/dS>1 shown in Table 4, with markedly higher average expression breadth (2.5 fold) for ovary-biased_TSSB_ (6.74±0.74) than for testis-biased_TSSB_ (2.73±0.72) genes (MWU-test P<0.05, Table 4). These patterns observed using whole-gene dN/dS values in this cricket system provide empirical data consistent with the theoretical proposition that that the fewer tissues a gene is expressed in, the more its adaptive evolutionary potential may be enhanced, likely by relaxing purifying selection imposed by multiple cross-tissue functions (Otto, 2004; Larracuente et al., 2008; Mank et al., 2008; Mank & Ellegren, 2009; Meisel, 2011). Our data thus specifically suggest that this hypothesis can apply to sex-biased genes (Mank & Ellegren, 2009). We note nonetheless that given the close relatedness between the two *Gryllus* species studied here, this might potentially elevate the overall genome-wide dN/dS including the portion with dN/dS> 1 ((Mugal et al., 2014) (see below section “*3.8 Close relatedness of Gryllus taxa*”), and thus further studies of dN.dS using additional *Gryllus* species as data becomes available will help test the rigor of these patterns across the genus.

**Table 4.**
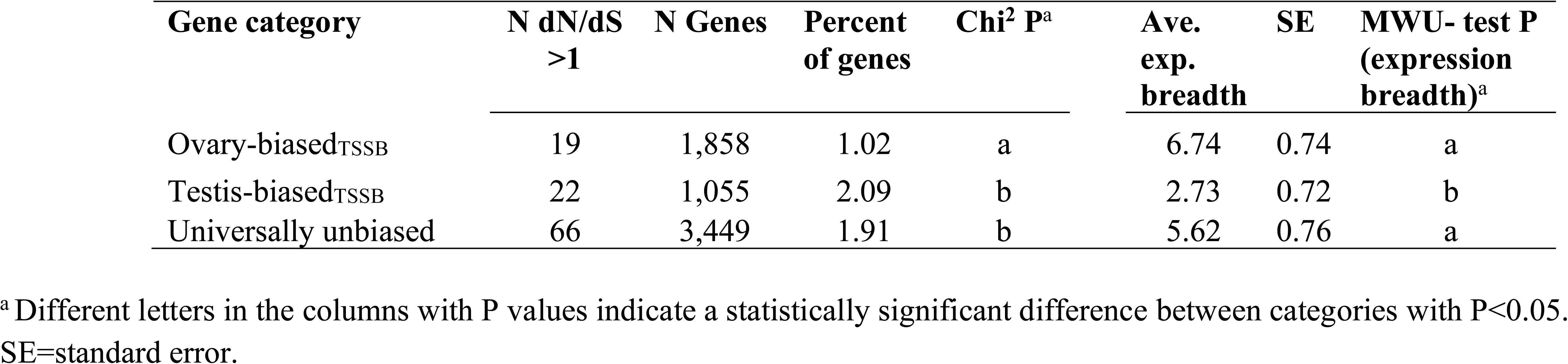
The proportion of genes with sex-biased_TSSB_ gonadal and universally unbiased expression in *G. bimaculatus* that had dN/dS>1, and their expression breadth across tissues (average number of nine tissues with expression >5 FPKM).

| Gene category | N dN/dS<br>>1 | N Genes | Percent<br>of genes | Chi<sup>2</sup> P<sup>a</sup> | Ave.<br>exp.<br>breadth | SE | MWU- test P<br>(expression<br>breadth)<sup>a</sup> |
| --- | --- | --- | --- | --- | --- | --- | --- |
| Ovary-biased<sub>TSSB</sub> | 19 | 1,858 | 1.02 | a | 6.74 | 0.74 | a |
| Testis-biased<sub>TSSB</sub> | 22 | 1,055 | 2.09 | b | 2.73 | 0.72 | b |
| Universally unbiased | 66 | 3,449 | 1.91 | b | 5.62 | 0.76 | a |

<sup>a</sup> Different letters in the columns with P values indicate a statistically significant difference between categories with P<0.05. SE=standard error.

We further assessed whether there was evidence of positive selection for sex-biased brain genes, which were much less common than those from the gonad (Table S3, Fig. 2). The only gene with whole-gene dN/dS >1 (=3.675, GBI_19557-RB, Table 1) was of unknown function and was expressed primarily in the male brain (number tissues with >5 FPKM =1 tissue). Thus, this result is also concordant with adaptive evolution facilitated by low pleiotropy. The female-biased brain gene with the highest dN/dS of 0.9735 matched *D. melanogaster kekkon3*. This value (near one) could suggest a history of neutral evolution, but may also reflect positive selection at multiple codon sites in that gene; we cannot distinguish between these two possibilities using gene-wide dN/dS.

As a follow-up supplemental analysis to gene-wide dN/dS, we examined positive selection among species at specific codon sites using branch-site analysis (with *G. bimaculatus* as the target branch) (Yang, 2007), based on three-way alignments of *G. bimaculatus*, *G. assimilis* and an available cricket outgroup species *Laupala kohalensis* (Blankers et al., 2018; Ylla et al., 2021). The results are described in Text File S1 and Table S8. It should be emphasized the assessment is inherently very conservative given it only includes the subset of genes with high confidence three-way reciprocal orthologs among the three species (that is, only 26,7% of the 7,220 genes with orthologs in the two *Gryllus* species had three-species orthologs, see Materials and Methods, and Text File S1). Nonetheless, we found that a non-negligible portion of the male- and female-biased_TSSB_ gonadal genes showed positive selection (≥9.6%), and that only minor variation was observed between groups, perhaps due to the conserved nature of the analysis (Table S8). Three sex-biased brain genes that were studied in Table 1 (among ten of the 65 in Table 1 that had three-species orthologs available for analysis, Table S8) showed positive selection using branch-site analysis (GBI_05906-RA, GBI_09477-RB, GBI_05452-RB, Table S8). This result is consistent with the hypothesis of a history of adaptive evolution in the brain, possibly elevating dN/dS (Fig. 3AB).

It is worth noting that for the branch-site analysis, we found that a small subset of *G. bimaculatus* genes that were female-biased in the somatic reproductive system (six of 33 genes (18.2%) with three-species orthologs), which includes the reproductive tract and/or spermathecae, tended to evolve adaptively using branch-site analysis (Table S8). In this context, the result suggests that a small number of female-biased reproductive system genes may evolve adaptively, potentially in response to sexual selection pressures, as suggested in flies (Swanson et al., 2004; Prokupek et al., 2008), in this cricket taxon. Further studies using more powerful branch-site positive selection tests (Yang, 2007) as genomic data emerge in even more crickets, and/or population genetics analysis of frequencies of codon mutations (McDonald & Kreitman, 1991), will further reveal the scale of positive selection at specific codon sites in the sex-biased genes from various tissues. Such analyses will also allow further evaluation of the link between positive selection (dN/dS>1) and gene pleiotropy that was suggested for gonads using the gene-wide dN/dS herein (Table 4, Fig. 4), and permit additional evaluation of this relationship for the brain, which had relatively few sex-biased genes with which to consider this specific relationship (of dN/dS>1 and pleiotropy) using gene-wide dN/dS (Fig. 2, Table 1).

### 3.8 Close relatedness of *Gryllus* taxa

The study of closely related species such as *G. bimaculatus* and *G. assimilis* as conducted herein allows for examination of genes with unsaturated substitutions and thus accurate measures of dN/dS (see section “*3.2.1 Rates of evolution*” for median dN and dS values), as applied in other studies within insect genera (Zhang et al., 2007; Baines et al., 2008; Meisel, 2011; Assis et al., 2012; Whittle & Extavour, 2017; Jaquiery et al., 2018). We note that very close relationships have been proposed in theory for some unicellular and viral systems (Rocha et al., 2006; Kryazhimskiy & Plotkin, 2008), and possibly some multicellular eukaryotes (Mugal et al., 2020), to potentially affect dN/dS due to a short time periods to fix or remove polymorphic mutations (see also counterevidence from (Gibson & Eyre-Walker, 2019)). In the present study, we propose that our core results are apt to be minimally influenced by any potential such effect, given that all genomic analyses were conducted in an identical manner for sex-biased genes from all tissues and for the same two species, and thus the time of divergence is the same throughout the two genomes. Nonetheless, follow-up studies in more species of *Gryllus* should consider the degree of relatedness in potentially shaping dN/dS among taxa. Further, the combined analyses of interspecies dN/dS data with polymorphism-level genomics data will allow discernment of whether any degree of nonsynonymous mutations may remain polymorphic (yet unfixed) between closely related cricket species. Unlike widely studied insects such as *Drosophila* that have vast available polymorphism and species genomic datasets (Wang et al., 2015; Gramates et al., 2017), studies testing hypotheses on the relationship between time since divergence and dN/dS in *Gryllus* will become feasible as more species genomes, as well as genome-wide population level datasets in multiple species, become available in the future.

## 4 CONCLUSIONS

Here we have conducted comprehensive assessment of sex-biased gene expression in reproductive and nervous system tissues, and revealed their relationships to potential pressures on protein sequence evolution, in a cricket model system. We have demonstrated the consistent tendency for rapid evolution of sex-biased brain genes, particularly female-biased brain genes, (Fig. 3, Table 1), and of male-biased genes from the gonad, in *G. bimaculatus*. Further, our data suggest a direct link between low pleiotropy and elevated dN/dS of sex-biased genes in the brain and the gonad (Fig. 3, Fig. 4) that may reflect relaxed purifying selection, which in turn may permit elevated instances of positive selection (Table 4) (Otto, 2004; Larracuente et al., 2008; Mank et al., 2008; Mank & Ellegren, 2009; Meisel, 2011). We speculate that the features of this cricket’s mating biology may give rise to sexual selection and thus contribute at least partly towards the accelerated evolution of the sex-biased brain genes, and male-biased gonadal genes, in this taxon.

Suggested significant directions for future studies include the following approaches: First, research on sex-biased gene expression from different brain regions may further decipher its relationship to protein evolution (Tuller et al., 2008), and the possible roles of allometric scaling (Montgomery & Mank, 2016). Second, investigation of the involvement of sex-biased brain genes in gene pathways and networks, and their expression breadth across even more tissue types than those studied herein, may help elucidate why they often evolve rapidly. Third, similar studies as conducted herein in more divergent *Gryllus* species and in other genera such as *Drosophila* may help reveal whether the relationships between sex-biased expression and dN/dS varies over evolutionary time (Mugal et al., 2014), and/or is affected by the turnover in sex-biased expression status (Zhang et al., 2007; Whittle & Extavour, 2019). Fourth, additional studies should consider potential differences in sex-biased expression of alternately spliced mRNAs among taxa, as high confidence genome-wide splicing variants are further refined for the recent *G. bimaculatus* genome (Ylla et al., 2021) and as whole genome and large-scale RNA-seq data (allowing splicing predictions) emerge in other comparable *Gryllus* species, some variants of which may be involved in sexual differentiation (Nagoshi et al., 1988; Wexler et al., 2019). Fifth, refinement of the *G. bimaculatus* genome to discern the sex chromosomes and autosomes and gene localizations, combined with expression data, will allow further testing of any putative role of dosage compensation and faster-X effect on rapid evolution of sex-biased genes from the brain and gonad (Parsch & Ellegren, 2013; Whittle et al., 2020). Sixth, the sequencing of additional *Gryllus* genomes and/or generation of population sequence data for *G. bimaculatus* may allow MacDonald-Kreitman tests and more powerful positive branch-site selection tests (McDonald & Kreitman, 1991; Yang, 2007) than available herein, particularly for those with small sample sizes of sex-biased genes such as the brain. Seventh, assessment of sex-biased gene expression in *G. bimaculatus* adult males and females should be conducted in a courtship environment with male-male rivals, and/or with multiple females exposed to few males (female-female competition), and include assessments of the putative roles of acoustics-related genes (*cf.* (Kasumovic et al., 2016; Pascoal et al., 2018; Pascoal et al., 2020). Given that mating behaviors may be largely mediated by gene expression in male and female brains in *Gryllus* (Matsumoto & Sakai, 2000; Haberkern & Hedwig, 2016; Sakai et al., 2017), and in other insects such as *Drosophila* (Fowler et al., 2019), such follow-up research in the brain will be valuable to better understand the potential ties between mating behaviors, sex-biased expression, and protein sequence evolution. Finally, the study of sex-biased expression in brain and gonad among insects that have known differences in their mating biology (including for example variation in testis size, sperm mixing, degree of female-female competition, mate choice (cf. Harrison et al., 2015)), including among additional species of *Gryllus*, will help further decipher whether and how protein sequence evolutionary rates may be shaped by these various mechanisms of sexual selection across a phylogeny.

## Acknowledgements

The authors thank Dr. Guillem Ylla for providing early access to the assembled *G. bimaculatus* and *L. kohalensis* genomes and members of the Extavour lab for discussions. The services of the Bauer core sequencing facility at Harvard University are appreciated. We also thank the anonymous reviewers for valuable comments that helped improve our manuscript.

## Author contributions

CAW, AK and CGE designed the study. AK reared G. *bimaculatus* and *G. assimilis* and sampled tissues for RNA-seq. CAW analyzed the data and wrote the manuscript with contributions by AK and CGE. All authors read and approved the final manuscript.

## Text File S1

### Assembly of *G. assimilis* RNA-seq data

To assess dN/dS, we compared the annotated genes in *G. bimaculatus* to the CDS list generated for *G. assimilis*. Applying Trinity and PlantTribes (see Methods) to the trimmed reads in Table S2, we obtained 33,089 non-redundant transcripts with a median and mean length of 540 bp and 784.3 bp respectively (standard error=30.3). The BUSCO score (Seppey et al., 2019) to the Arthropoda conserved gene set of 1,066 genes, showed 86.7% CDS had complete sequence matches, 8.6% were fragmented matches, and 4.7% were missing. The latter may represent gene losses in this species, and/or genes excluded from the assembly. Thus, this suggests high efficiency of the assembly spanning a major portion of arthropod genes. From this list, we used ORF predictor with *G. bimaculatus* CDS as a reference and BLASTX to identify *G. assimilis* CDS. We found 25,128 CDS (including isoforms) with a start codon and no unknown or ambiguous nucleotides, which were used for analyses. Reciprocal BLASTX of the 15,539 *G. bimaculatus* CDS to the *G. assimilis* CDS yielded 7,919 putative orthologs between the two species (e<10^-6^ in both forward and reverse matches). Retaining only those putative ortholog matches that after alignment had both dN and dS values <1.5, and thus were unsaturated, yielded a total of 7,220 high confidence between-species orthologs that were used for all our dN/dS analyses.

### Comparison of sex-biased gonadal expression in *G. bimaculatus* and *G. assimilis*

As described in our main text, our core target for expression analysis was *G. bimaculatus*, which has an assembled genome with complete or near complete CDS (Ylla et al., 2021), and *G. assimilis* was used primarily as a reference for assessment of protein divergence. Nonetheless, we assessed the degree of conservation of gene expression between species for the 7,220 genes with orthologs for the gonads (largest N values of all tissues, Table S3) between these two species. The results showed that gene expression in *G. assimilis* gonads was strongly correlated to that in *G. bimaculatu*s, with Spearman’s R=0.780 and 0.775 (P<2X10^-7^) for ovary and testis expression respectively (Fig. S2AB). In addition, 65.9 and 65.8% of all gonadally expressed genes (among the 7,220 with orthologs) that were defined as female- and male-biased in *G. bimaculatus* (ALL genes sex-biased in testis regardless of status in other tissues, N=2,043 and 1,225 respectively) had the same status in *G. assimilis*. This suggests substantial turnover in sex-biased status, a pattern observed for gonadal tissues in studied species of *Drosophila* (Zhang et al., 2007; Assis et al., 2012; Harrison et al., 2015; Whittle & Extavour, 2019b). Importantly, for genes with the same (conserved) sex-biased status in the two species, dN/dS was highest in testis-biased genes (median=0.127) and lower in unbiased (0.114), and ovary-biased (0.097) genes (MWU-tests P<0.05 for all paired contrasts) (Fig. S2C). Moreover, genes that were testis-biased in only one species (either *G. bimaculatus* or *G. assimilis*) and unbiased in the other species had elevated dN/dS values as compared to their ovary-biased counterparts (MWU-test P<0.05 for each contrast, Fig. S2D). Thus, the accelerated evolution of testis-biased genes is robust to whether the sex-biased status is observed in one species, or both species, in this taxon. All our remaining analysis is using sex-biased genes from our annotated model *G. bimaculatus*.

### Assessment of expression in male accessory glands and seminal fluid proteins

We considered the evolution of genes specifically linked to the male accessory glands in *G. bimaculatus*, including those defined as putative orthologs to *D. melanogaster* seminal fluid proteins (SFPs; see below paragraph (Sepil et al., 2019)). First, we took a broad approach to study all male accessory gland-specific genes identified using our RNA-seq dataset (Table S1). Prior study of two species of crickets (*G. firmus* and *G. pennsylvanicus*) identified transcripts from the male accessory glands or SFPs, whereby some were suggested to evolve rapidly (Andres et al., 2006; Andres et al., 2013). Herein, we have the advantages of large-scale RNA-seq data from multiple tissue types, and an annotated *G. bimaculatus* genome (N=15,539 CDS) (Ylla et al., 2021), to identify male-accessory gland-specific genes in this species. We report a total of 30 genes expressed in the male accessory glands with no expression (0 FPKM) in all eight other studied male and female tissues (Table S1).

Functional predictions of the 30 male accessory gland-specific genes using *D. melanogaster* orthologs (Table S6, e<0.001, see Methods) revealed seven genes with a match. Two of these *G. bimaculatus* genes are predicted orthologs of *painless* and *Sox100B*, which have functions in male reproduction in *D. melanogaster*; the former is involved in courtship and olfactory signalling (Table S6). Both genes were expressed at low levels (FPKM<1) in male accessory glands in *G. bimaculatus*. Only one of the 30 accessory gland specific genes had a match in the two *Gryllus* species (3.33.%, Table S6, which had very strict match criteria, see Methods). Several of the *G. bimaculatus* accessory gland genes with no *G assimilis* or *D. melanogaster* ortholog matches had relatively high expression levels (e.g., 16 to 347 FPKM; Table S6), and we speculate they could comprise orphan genes that have evolved essential male sexual functions specifically in *G. bimaculatus* (Tautz & Domazet-Loso, 2011; Whittle & Extavour, 2019a). Overall, the nearly complete lack of high confidence orthologs between *G. bimaculatus* and *G. assimilis* suggests there has been rapid evolution of male accessory gland specific genes resulting in similarity too low for ortholog detection using these methods. Alternatively, these results may reflect a history of some lineage-specific gene losses or gains of these rapidly changing genes (Haerty et al., 2007; Tautz & Domazet-Loso, 2011).

#### Seminal fluid proteins

Seminal fluid proteins (SFPs) play significant roles in sperm vitality, sperm storage in the female spermatheca after mating, and in fertilization (Sepil et al., 2019). In studied systems to date, which have preferentially focused on primates and *Drosophila*, genes described as SFPs have been found to evolve rapidly and/or adaptively (Swanson et al., 2001; Clark & Swanson, 2005; Haerty et al., 2007). While it may be predicted that rapid evolution of SFPs might be more pronounced in systems where females have multiple mates (such as *G. bimaculatus*) than those that are monogamous, this expected pattern was not observed for a study of 18 candidate SFPs in butterflies, where monogamy was unexpectedly linked to fast evolution of SFPs, perhaps due to relaxed selective constraints (Torgerson et al., 2002; Walters & Harrison, 2011). Research on SFPs in more diverse insects with well-described mating biology are thus needed (Walters & Harrison, 2011). *G. bimaculatus* has high female polyandry, complete sperm mixing, and exhibits extensive pre- and post-mating female choice (Simmons, 1986; Morrow & Gage, 2001). Using the recently available list of 134 SFPs in *D. melanogaster* (shown in Table S7, (Sepil et al., 2019)), we found that only 20 genes had identifiable putative orthologs in *G. bimaculatus* genes (14.9%). This is much lower than the 64.5% genome-wide rate of putative ortholog detection between these two species (Chi-square with Yates’s correction P<0.001). Thus, the lack of putative SFP orthologs is consistent with especially rapid evolution (Haerty et al., 2007; Tautz & Domazet-Loso, 2011) of the SFP genes following the divergence of the lineages leading to *D. melanogaster* and *G. bimaculatus*.

Among the 20 putative *G. bimaculatus* SFP genes, seven were included among the subset of 7,220 genes with between-species orthologs in *Gryllus* (Table 4; note that none of these were among the 30 accessory gland-specific genes reported above). It has been inferred that SFPs tend be produced in insect accessory glands, as well as in the testis or male somatic reproductive system tissues (Sepil et al., 2019). Indeed, we found that each of these seven putative *Gryllus* SFPs exhibited expression within the testis, male somatic reproductive system, and the male accessory glands (between 0.2 to 1392.5 FPKM depending on tissue, with one exception, testis for GBI_14450-RA FPKM=0, Table 4). Significantly, for each of these seven putative cricket SFPs, we also found that the dN/dS values were consistently well above the median observed for all studied genes in the genome (which was 0.115 across all 7,220 genes, shown in Fig. 3A). Specifically, the values were 0.149 (*Phm*), 0.220 (*Npc2b*), 0.230 (*Ggt-1*), 0.250 (*regucalcin*), 0.287 (*Spn28F),* 0.344 (*Spn38F*) and 2.48 (*Obp56g*) (Table 4). Thus, the putative SFPs in the crickets studied here have evolved very rapidly, a feature shared with the SFPs that have been studied in the fellow insect *D. melanogaster* (Haerty et al., 2007; Sepil et al., 2019). It should be noted that while we consider it unlikely, we cannot exclude the possibility that some accessory gland or SFP CDS may be expressed at extremely low levels in the *G. assimilis* tissue types used for RNA-seq, causing an apparent absence of orthologs to *G. bimaculatus* in that assembly. However, we consider this unlikely given the number of tissues we assessed, including the male accessory glands (Table S2). Moreover, this would not explain the apparent paucity of *G. bimaculatus* SFP orthologs relative to those in the *D. melanogaster* genome. Thus, we suggest the absence is best explained by rapid divergence that obscures ortholog detection, and/or from gene losses or gains (Haerty et al., 2007; Tautz & Domazet-Loso, 2011).

A role of positive selection for at least one SFP gene in *Gryllus* is supported by the fact that the dN/dS value was >1 (was 2.5, Table 4) for the odorant binding SFP protein *Obp56g*. In *D. melanogaster*, *Obp56g* was first recognized as an SFP using proteomics of seminal fluid in mated females (Findlay et al., 2008), was later affirmed as a protein stored in male reproductive tissues (Takemori & Yamamoto, 2009) (which we have confirmed also express this gene in crickets: (Table 4)), and was stringently verified as an SFP by Sepil and colleagues (2019).

### Branch-site analysis for *G. bimaculatus*

Three-way reciprocal BLASTX of *Laupala kohalensis* to *G. bimaculatus* and *G. assimillus* yielded 4,523 genes with putative orthologs. Using free-ratio branch analyses of the three species, we found dN was largely unsaturated for the *L. kohalensis* branch, with a median of 0.10. However, dS values were particularly high (median=3.3), suggesting a high mutation rate in this organism. Including only genes with dN and dS <3 yielded 1,933 genes with confidence orthologs in *L. kohalensis* (26.7% of the 7,220 genes with *G. bimaculatus* and *G. assimillus* orthologs). This conservative approach favors study of the slowly evolving genes in each sex-biased category. We found instances of positive selection at specific sites in the *G. bimaculatus* branch for sex-biased genes from all studied tissue types (2XlnL P <0.05, Table S8). For instance, we found 11.8%, 9.6% and 10.9% of studied genes exhibited positive selection for ovary-biased_TSSB_, testis-biased_TSSB_ and universally unbiased genes (2XlnL P per gene <0.05; Table S8). The use of conserved genes, however, biases these testis-biased estimates of positive section downward (as fast evolving genes are excluded more often: 23.7% of testis-biased genes had three-way orthologs, versus 29.8% for ovary-biased genes). Further, while the number of genes, and thus three-way orthologs, were uncommon outside the gonads (N=4-33 depending on tissue; Table S8), we found that more than three times as many female-biased than male-biased somatic reproductive system genes exhibited branch-site selection (18.2% versus 5%; but this was not statistically significant, Chi-square P=0.17, Table S8), suggesting that this narrowed level of analysis (branch-site analysis of conserved genes), may concur with the notion that some genes from the female reproductive tract and/or spermathecae, which store sperm after mating, tend to evolve adaptively due to sexual selection pressures (Swanson et al., 2004; Prokupek et al., 2008). Future studies using more closely related cricket genomes as data emerge will be needed to enhance the power of detecting branch-site positive selection using branch-site analysis.

## SUPPORTING INFORMATION

**Fig. S1.**
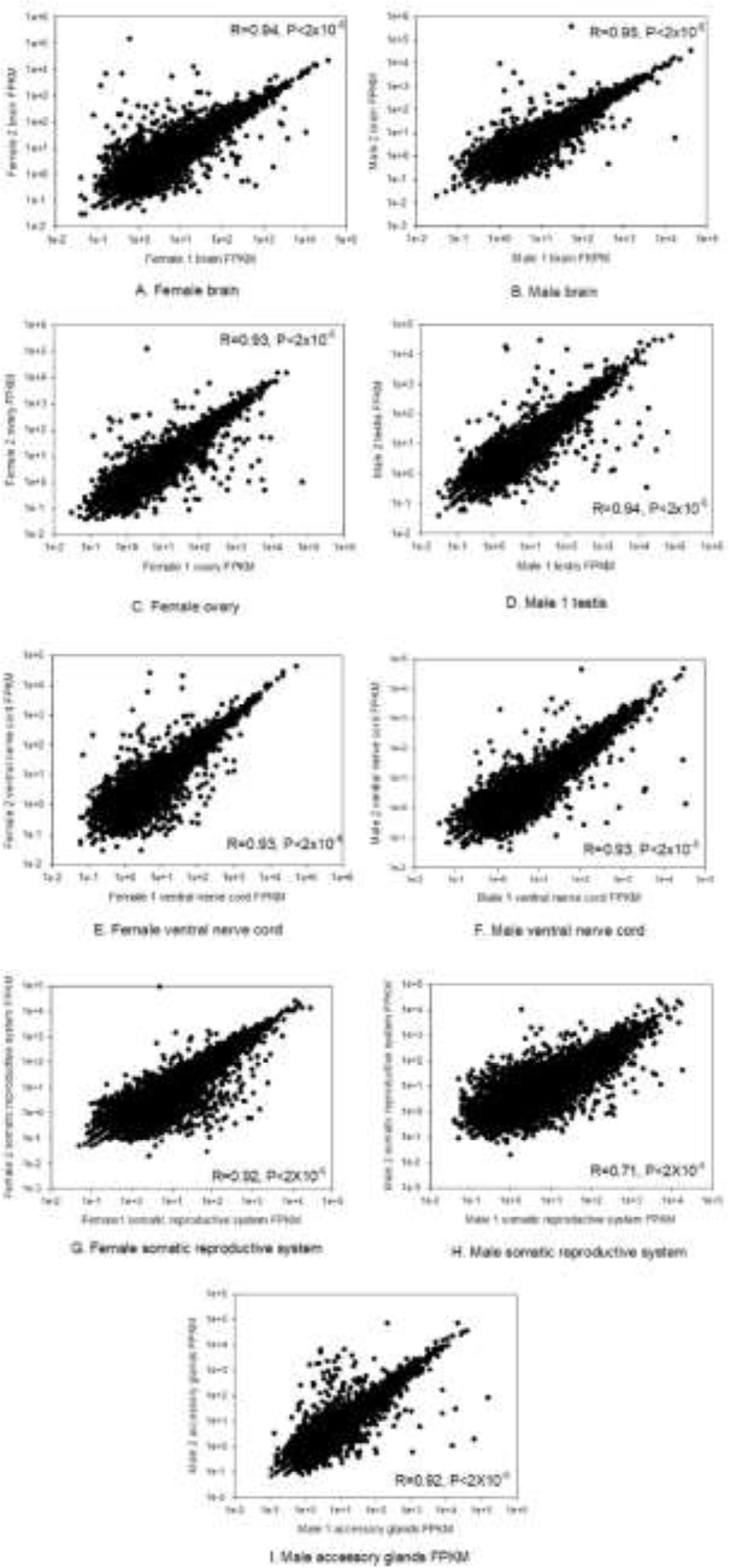
The Spearman correlation (R) in FPKM across all all 15,539 genes in *G. bimaculatus* for each of the tissues under study. A) female brain; B) male brain; C) ovary; D) testis; E) female ventral nerve cord; F) male ventral nerve cord; G) female somatic reproductive system; H) male somatic reproductive system; I) male accessory glands.

**Fig. S2.**
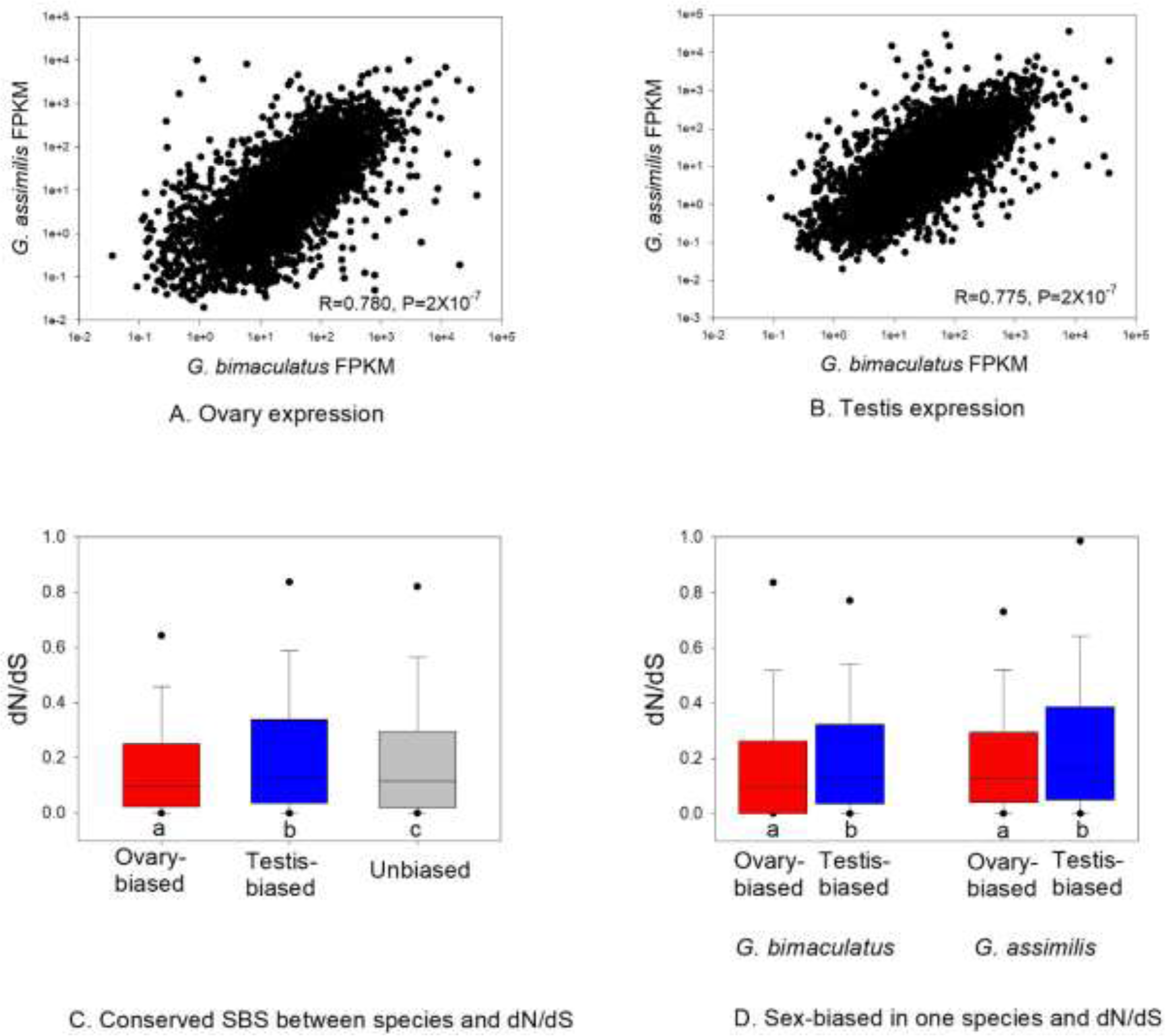
Expression of genes in *G. bimaculatus* and *G. assimilis* for A) ovaries and B) testes (Spearman’s R and P are shown; expression per gene is the average across samples per species). C) Box plots of dN/dS for genes that have the same sex-biased status (SBS) in both *G. bimaculatus* and *G. assimilis* and; D) Box plots of dN/dS for genes that are ovary-biased or testis-biased in only one species. Different letters under bars in C and each pair of bars in D indicate MWU-P<0.05.

**Table S1.** The RNA-seq datasets for each of the male and female tissue types under study for *G. bimaculatus*. The number of reads (single-end) before and after trimming with BBduk (https://jgi.doe.gov/data-and-tools/bbtools/) is shown. The data are available at the Short Read Archive (SRA) under the project identifier PRJNA564136.

| Sample | Tissue | Sample Name | No. Reads |  |
| --- | --- | --- | --- | --- |
|  |  |  | Before trimming | After trimming |
| Male 1 | Accessory gland | AK-28_S6.R1 | 8,519,999 | 8,455,381 |
|  | Brain | AK-25_S3.R1 | 10,927,264 | 10,543,501 |
|  | Somatic reproductive system | SHC-18_S14.R1 | 32,497,283 | 32,430,843 |
|  | Testes | SHC-17_S13.R1 | 19,928,912 | 19,751,731 |
|  | Ventral nerve cord | AK-26_S4.R1 | 11,488,521 | 11,140,299 |
| Male 2 | Accessory gland | AK-35_S13.R1 | 15,110,718 | 14,973,668 |
|  | Brain | AK-32_S10.R1 | 18,039,328 | 17,850,399 |
|  | Somatic reproductive system | AK-31_S9.R1 | 11,993,680 | 11,702,596 |
|  | Testes | AK-30_S8.R1 | 13,672,147 | 13,529,248 |
|  | Ventral nerve cord | AK-33_S11.R1 | 11,677,747 | 11,445,159 |
| Female 1 | Brain | AK-39_S17.R1 | 13,920,966 | 13,750,206 |
|  | Ovary | AK-37_S15.R1 | 21,725,208 | 21,128,416 |
|  | Somatic reproductive system | AK-38_S16.R1 | 13,870,827 | 13,718,497 |
|  | Ventral nerve cord | AK-40_S18.R1 | 12,599,661 | 12,341,413 |
| Female 2 | Brain | AK-45_S23.R1 | 19,312,301 | 19,036,974 |
|  | Ovary | AK-43_S21.R1 | 27,627,122 | 27,049,583 |
|  | Somatic reproductive system | AK-44_S22.R1 | 11,688,814 | 11,539,571 |
|  | Ventral nerve cord | AK-46_S24.R1 | 13,591,143 | 13,143,568 |

**Table S2.** The RNA-seq datasets for each of the male and female tissue types under study for *G. assimilis*. The number of reads (single-end) before and after trimming with BBduk is shown. All tissue samples were used for the RNA-seq assembly, and * indicates those samples used for gonadal expression (Exp.) analyses for comparison to *G. bimaculatus* tissue samples. The data are available at the Short Read Archive (SRA) under the project identifier PRJNA564136.

| Sample | Tissue<sup>a</sup> | File Name | No. Reads |  | Exp. |
| --- | --- | --- | --- | --- | --- |
|  |  |  | Before trimming | After trimming |  |
| **Male 1** | Accessory gland | AK_1-6_S6_R1_001 | 7,232,269 | 6,927,036 |  |
|  | Brain | AK_1-3_S3_R1_001 | 414,137 | 397,903 |  |
|  | Carcass | AK_1-5_S5_R1_001 | 12,162,122 | 11,909,287 |  |
|  | Somatic reproductive system | AK_1-2_S2_R1_001 | 9,786,892 | 9,218,317 |  |
|  | Testes | AK_1-1_S1_R1_001 | 15,489,005 | 14,677,775 |  |
|  | Ventral nerve cord | AK_1-4_S4_R1_001 | 8,085,561 | 7,778,520 |  |
| **Male 2** | Accessory gland | AK_1-12_S12_R1_001 | 9,684,938 | 9,395,491 |  |
|  | Brain | AK_1-9_S9_R1_001 | 10,251,515 | 9,935,844 |  |
|  | Carcass | AK_1-11_S11_R1_001 | 13,140,136 | 12,739,689 |  |
|  | Somatic reproductive system | AK_1-8_S8_R1_001 | 9,038,153 | 8,700,106 |  |
|  | Testes | AK_1-7_S7_R1_001 | 15,015,495 | 14,252,214 | * |
|  | Ventral nerve cord | AK_1-10_S10_R1_001 | 159,249,632 | 153,314,813<sup>b</sup> |  |
| **Male 3** | Accessory gland | AK_1-18_S18_R1_001 | 9,301,332 | 8,771,926 |  |
|  | Brain | AK_1-15_S15_R1_001 | 8,556,018 | 8,269,869 |  |
|  | Carcass | AK_1-17_S17_R1_001 | 11,258,971 | 11,016,352 |  |
|  | Somatic reproductive system | AK_1-14_S14_R1_001 | 11,468,584 | 11,055,961 |  |
|  | Testes | AK_1-13_S13_R1_001 | 11,082,626 | 10,637,566 | * |
|  | Ventral nerve cord | AK_1-16_S16_R1_001 | 9,501,325 | 9,141,580 |  |
| **Female 1** | Brain | AK_1-21_S21_R1_001 | 12,314,902 | 11,529,951 |  |
|  | Carcass | AK_1-23_S23_R1_001 | 10,318,471 | 9,965,655 |  |
|  | Ovary | AK_1-19_S19_R1_001 | 12,968,675 | 12,330,995 | * |
|  | Somatic reproductive system | AK_1-20_S20_R1_001 | 20,180,007 | 19,613,713 |  |
|  | Ventral nerve cord | AK_1-22_S22_R1_001 | 13,818,212 | 13,322,784 |  |
| **Female 2** | Brain | AK_1-26_S26_R1_001 | 10,596,275 | 10,191,182 |  |
|  | Carcass | AK_1-28_S28_R1_001 | 9,471,179 | 8,987,504 |  |
|  | Ovary | AK_1-24_S24_R1_001 | 14,894,350 | 14,584,072 | * |
|  | Somatic reproductive system | AK_1-25_S25_R1_001 | 10,705,738 | 10,183,713 |  |
|  | Ventral nerve cord | AK_1-27_S27_R1_001 | 10,108,946 | 9,733,477 |  |
| **Female 3** | Brain | AK_1-31_S31_R1_001 | 9,543,257 | 9,388,801 |  |
|  | Carcass | AK_1-33_S33_R1_001 | 14,562,167 | 14,279,995 |  |
|  | Ovary | AK_1-29_S29_R1_001 | 10,900,114 | 10,725,546 |  |
|  | Somatic reproductive system | AK_1-30_S30_R1_001 | 9,846,659 | 9,641,174 |  |
|  | Ventral nerve cord | AK_1-32_S32_R1_001 | 7,954,359 | 7,795,480 |  |

<sup>a</sup> The carcass consists of body and head (minus the legs, wings, antennae, gut, Malpighian tubules, gonad and somatic reproductive system, ventral nerve cord, and brain); <sup>b</sup>The sample was divided into tenths and one-tenth of the reads were randomly used for assembly and expression analysis to approximately match the read number of other samples.

**Table S3.**
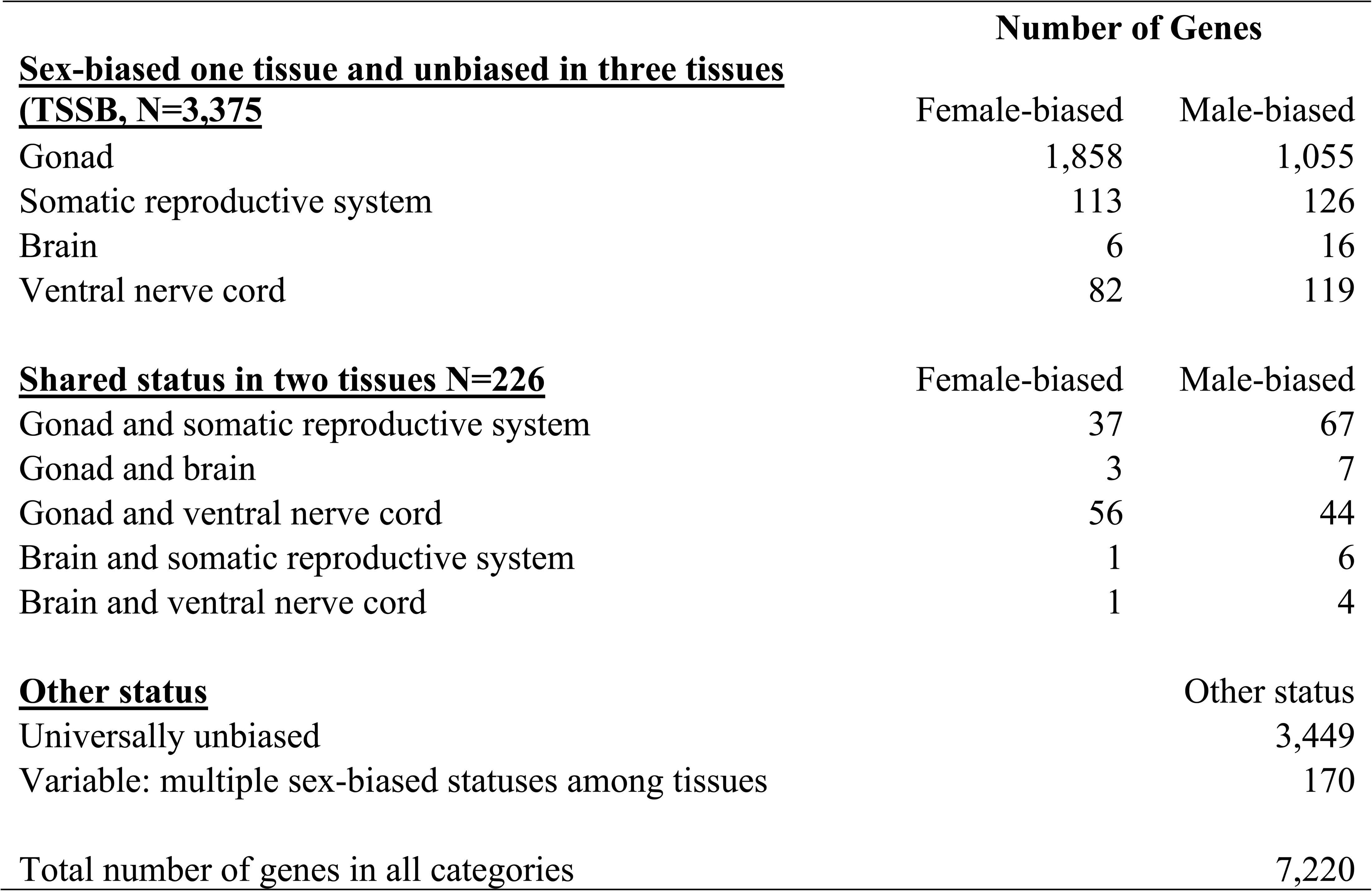
The number of genes in *G. bimaculatus* under study for dN/dS (those with between-species orthologs) that had female- or male-biased expression in one (of four) paired tissue types and were unbiased in expression for the three remaining tissue types (tissue-specific sex bias, TSSB). Genes with shared sex-biased status in two of four tissues (and unbiased in two) or other types of variation in SBS are also shown.

**Table S4.** The top GO functional classifications of genes with sex-biased brain expression (all genes regardless of interspecies orthologs, N values in Fig. 2). Annotation was determined in DAVID (Huang da et al., 2009) using *D. melanogaster* orthologs to *G. bimaculatus* genes. Functions are ranked by the percentage of genes matching its classification. P-values indicate the enrichment of genes involved in each function with lower values indicating greater enrichment.

| **Female-biased brain genes** | **Percent of Genes** | **P-value** |
| --- | --- | --- |
| Positive regulation of transcription from RNA polymerase II promoter | 12.1 | 1.10E-02 |
| Sensory perception of pain | 12.1 | 3.40E-02 |
| Transcription, DNA-templated | 12.1 | 4.20E-02 |
| **Male-biased brain genes** |  |  |
| Proteolysis | 11.3 | 2.90E-04 |
| Calcium ion binding | 8.5 | 6.70E-03 |
| Neuron remodeling | 4.2 | 2.10E-02 |

**Table S5.** The top GO functional classifications of genes with universally unbiased expression across all four paired male-female tissue types under study (those with interspecies orthologs for study). Annotation was determined in DAVID (Huang da et al., 2009) using *D. melanogaster* orthologs to *G. bimaculatus* genes. Functions are ranked by the percentage of genes matching its classification and those categories representing 5% or more of genes are shown. P-values indicate the enrichment of genes involved in each function with lower values indicating greater enrichment.

| GO keyword | Percent<br>of genes | P-value |
| --- | --- | --- |
| Coiled coil | 16.1 | 5.20E-16 |
| Hydrolase | 12.9 | 4.00E-03 |
| Transferase | 12.4 | 2.40E-21 |
| Nucleus | 11.2 | 8.30E-10 |
| Phosphoprotein | 11 | 3.40E-22 |
| Metal-binding | 10.8 | 6.00E-14 |
| Nucleotide-binding | 8.3 | 2.50E-15 |
| Alternative splicing | 8.1 | 4.40E-20 |
| Cytoplasm | 8 | 1.00E-17 |
| ATP-binding | 7 | 3.00E-16 |
| Developmental protein | 6.7 | 5.20E-11 |
| Zinc | 6.6 | 1.20E-09 |
| Transport | 5.1 | 1.70E-03 |
| Oxidoreductase | 5.0 | 4.30E-02 |

**Table S6.** The 30 genes that were expressed specifically in the male accessory glands and not in any of the eight other (male and female) tissue types in *G. bimaculatus*. The genes that had high confidence ortholog matches between *G. bimaculatus* and *G. assimilis* are shown (N=1, indicated by *), as well as those with putative orthologs identified in *D. melanogaster* (N=7), which had less strict criteria (for identification of a match for gene ontology purposes; See Methods). Gene functions were predicted using DAVID and *D. melanogaster* gene identifiers (Huang da et al., 2009).

| Row | *G. bimaculatus*<br>gene | *D. melanogaster*<br>match | Gene | GO putative functions | Expression<br>(FPKM) | dN/dS |
| --- | --- | --- | --- | --- | --- | --- |
| 1 | GBI_11980-RA* | FBgn0035154 | CG3344 | Peptidase S10, serine carboxypeptidase | 19.46 | 0.1039 |
| 2 | GBI_06110-RA | FBgn0035781 | CG8560 | proteolysis | 9.60 | - |
| 3 | GBI_00239-RA | FBgn0039084 | CG10175 | neuron cell-cell adhesion, synaptic transmission | 0.83 | - |
| 4 | GBI_14669-RA | FBgn0259215 | *Ionotropic receptor*<br>*93a (Ir93a)* | detection of chemical stimulus sensory perception | 0.70 | - |
| 5 | GBI_18443-RA | FBgn0035476 | CG12766 | oxidation-reduction process | 0.34 | - |
| 6 | GBI_06560-RA | FBgn0024288 | *Sox100B* | male gonad development | 0.27 | - |
| 7 | GBI_04401-RA | FBgn0060296 | *painless(pain)* | copulation, male courtship behavior, olfactory | 0.08 | - |
| 8 | GBI_08022-RA | No match | - | - | 347.25 | - |
| 9 | GBI_00292-RA | No match | - | - | 120.78 | - |
| 10 | GBI_13609-RA | No match | - | - | 41.80 | - |
| 11 | GBI_13241-RA | No match | - | - | 20.21 | - |
| 12 | GBI_14160-RA | No match | - | - | 16.75 | - |
| 13 | GBI_01608-RA | No match | - | - | 15.96 | - |
| 14 | GBI_05352-RA | No match | - | - | 8.13 | - |
| 15 | GBI_06938-RA | No match | - | - | 5.72 | - |
| 16 | GBI_21228-RA | No match | - | - | 5.29 | - |
| 17 | GBI_09340-RA | No match | - | - | 3.41 | - |
| 18 | GBI_17179-RA | No match | - | - | 3.39 | - |
| 19 | GBI_18175-RA | No match | - | - | 1.42 | - |
| 20 | GBI_03401-RA | No match | - | - | 1.21 | - |
| 21 | GBI_06890-RA | No match | - | - | 1.15 | - |
| 22 | GBI_06913-RA | No match | - | - | 1.08 | - |
| 23 | GBI_02473-RA | No match | - | - | 0.92 | - |
| 24 | GBI_02976-RA | No match | - | - | 0.74 | - |
| 25 | GBI_05550-RA | No match | - | - | 0.6 | - |
| 26 | GBI_00099-RA | No match | - | - | 0.49 | - |
| 27 | GBI_11938-RA | No match | - | - | 0.35 | - |
| 28 | GBI_16483-RA | No match | - | - | 0.35 | - |
| 29 | GBI_07340-RA | No match | - | - | 0.29 | - |
| 30 | GBI_04232-RA | No match | - | - | >0<sup>a</sup> | - |

<sup>a</sup> Highly variable FPKM among individuals and thus denoted as >0 FPKM.

**Table S7.**
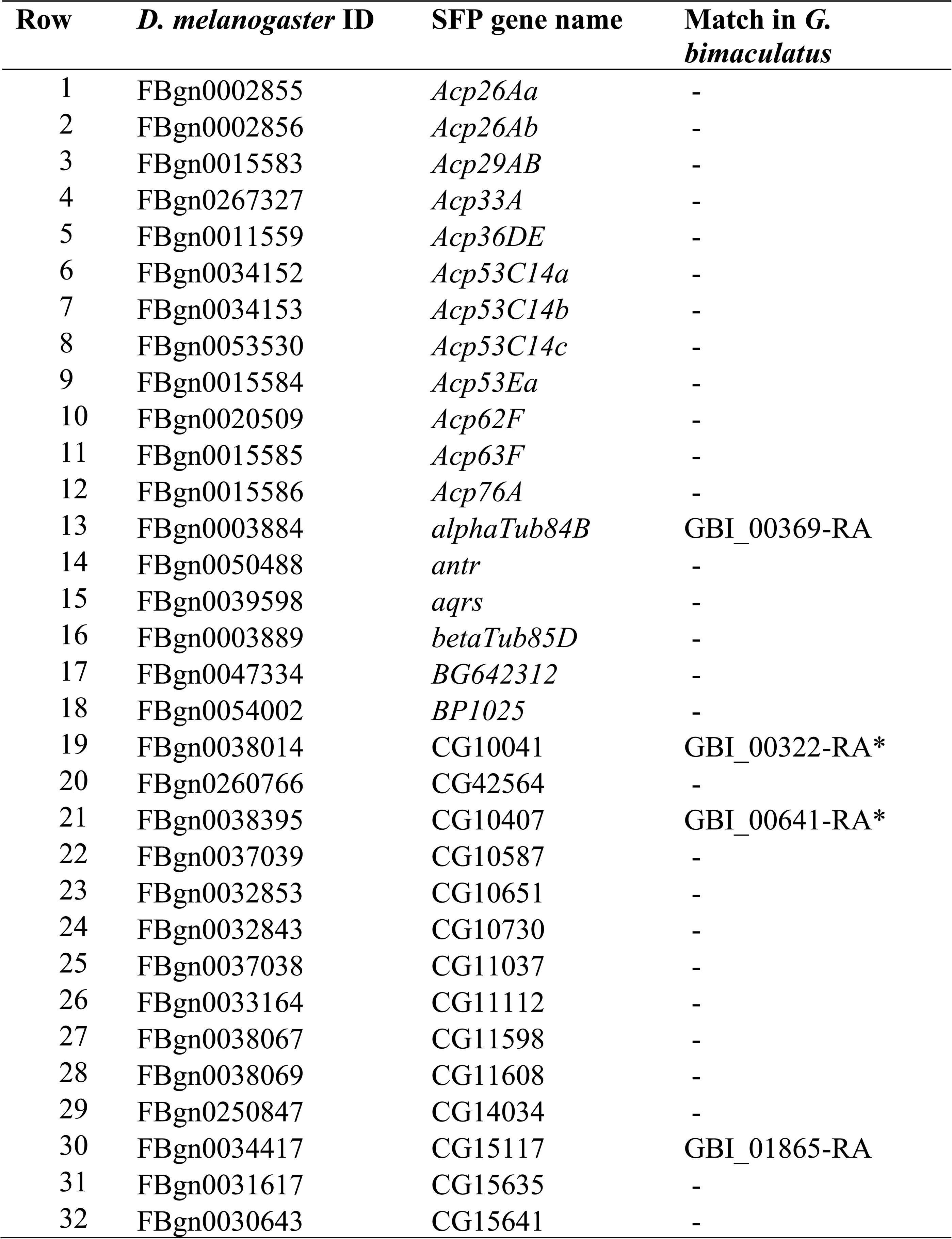
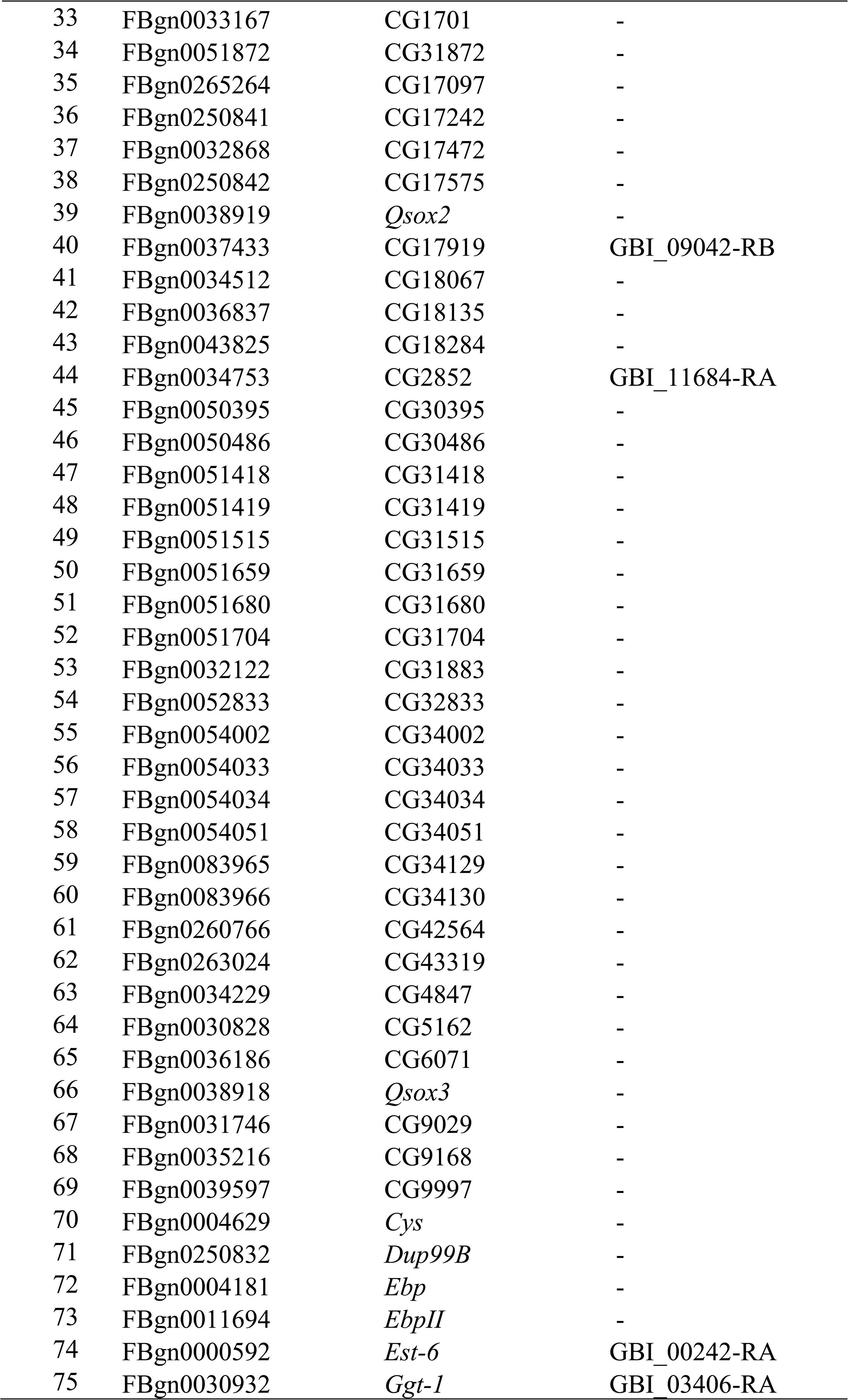
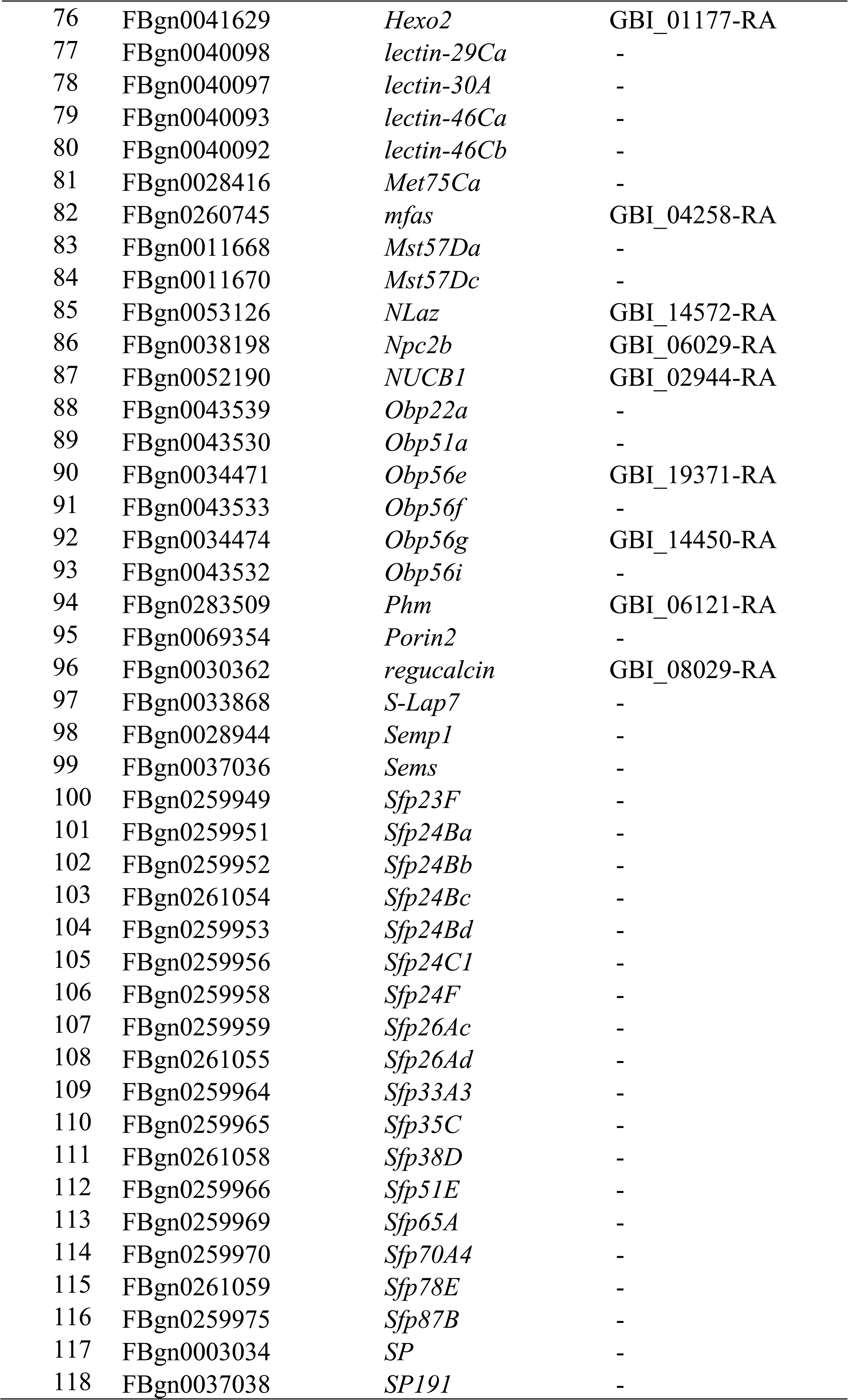
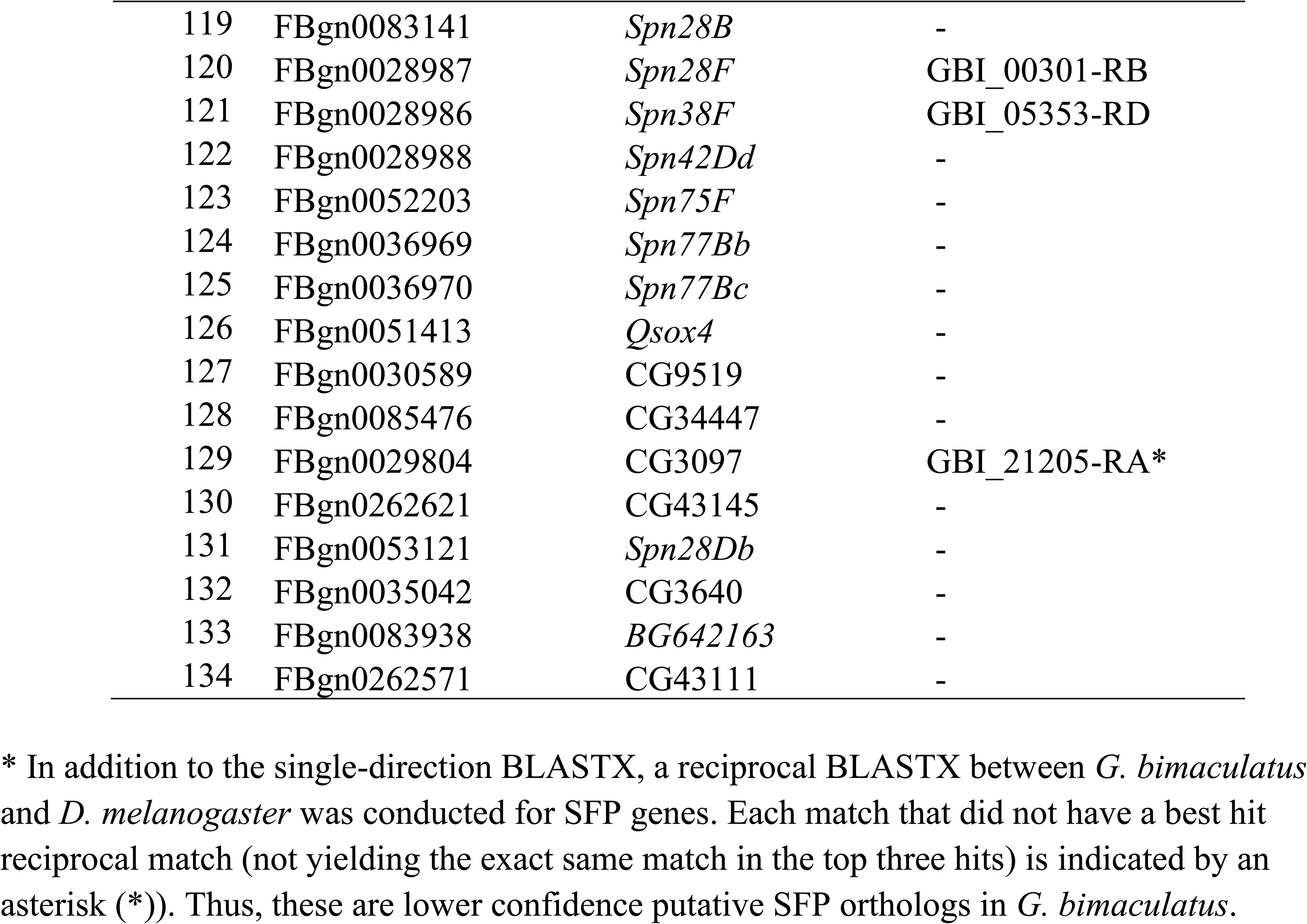
The 134 seminal fluid protein (SFPs) genes for the species *D. melanogaster* from Sepil et al. 2019 (Sepil et al., 2019) and their best BLASTX matches in the 15,539 genes of *G. bimaculatus*. Due to the extended phylogenetic distance between species, the list shows all putative orthologs identified using single forward BLASTX of *G. bimaculatus* to *D. melanogaster* using BLASTX (e<0.001). Results for those contained within the 7,220 genes with high confidence orthologs between *G. bimaculatus* and *G. assimilis* are shown in Table 4 within the main text.

**Table S8.** The number of orthologs identified between *G. bimaculatus* (GB) and *G. assimilis* (GA) and the outgroup *L. kohalensis* (LK). Among the orthologs studied for GB-GA paired analysis, the number of genes also having an LK ortholog after three-way reciprocal BLASTX and after excluding genes with dN or dS>3 are shown (designated as high confidence). Branch-site analysis results including 2XΔL are shown for all genes studied and for sex-biased TSSB genes (or all genes for the brain) and universally unbiased genes. Genes not belonging to any of these categories were excluded (Table S3).

|  | All<br>genes | Gonad-biased<sub>TSSB</sub> |  | Somatic reproductive<br>system-biased<sub>TSSB</sub> |  | Ventral-biased<sub>TSSB</sub> |  | Brain-biased<sub>ALL</sub> |  | Universally<br>unbiased |
| --- | --- | --- | --- | --- | --- | --- | --- | --- | --- | --- |
| <u>Identification of three-species orthologs</u> |  | Ovary-biased | Testis-biased | Female-biased | Male-biased | Female-biased | Male-biased | Female-biased | Male-biased |  |
| N GB-GA-LK putative orthologs (BLASTX) | 4,523<sup>a</sup> | 1,300 | 597 | 62 | 59 | 49 | 53 | 15 | 15 | 2,171 |
| N with dN and dS < 3 (High confidence) | 1,933<sup>a</sup> | 553 | 250 | 33 | 20 | 14 | 31 | 6 | 4 | 927 |
| <u>N branch-site test (2XΔlnL) P<0.05</u> | 220<sup>a</sup> | 65 | 24 | 6 | 1 | 1 | 7 | 1 | 2 | 101 |
| Percent of studied genes statistically significant | 11.38% | 11.75% | 9.60% | 18.18% | 5.00% | 7.14% | 22.58% | NA<sup>b</sup> | NA<sup>b</sup> | 10.90% |

<sup>a</sup>The total is for all genes with orthologs, including some not belonging to any of the sub-categories. <sup>b</sup>Too few genes were available to study for a reliable estimate.

